# The Wnt/TCF7L1 transcriptional repressor axis drives primitive endoderm formation by antagonizing naive and formative pluripotency

**DOI:** 10.1101/2022.05.18.492419

**Authors:** Paraskevi Athanasouli, Martina Balli, Anchel De Jaime-Soguero, Annekatrien Boel, Sofia Papanikolaou, Bernard K. van der Veer, Adrian Janiszewski, Annick Francis, Youssef El Laithy, Antonio Lo Nigro, Francesco Aulicino, Kian Peng Koh, Vincent Pasque, Maria Pia Cosma, Catherine Verfaille, An Zwijsen, Bjorn Heindryckx, Christoforos Nikolaou, Frederic Lluis

## Abstract

Early during preimplantation development and in heterogeneous mouse embryonic stem cells (mESC) culture, pluripotent cells are specified towards either the primed epiblast or the primitive endoderm (PE) lineage. Canonical Wnt signaling is crucial for safeguarding naive pluripotency and embryo implantation, yet the role and relevance of canonical Wnt inhibition during early mammalian development remains unknown. Here, we demonstrate that transcriptional repression exerted by Wnt/TCF7L1 promotes PE differentiation of mESCs and in preimplantation inner cell mass. Time-series RNA sequencing and promoter occupancy data reveal that TCF7L1 binds and represses genes encoding essential naive pluripotency factors and indispensable regulators of the formative pluripotency program, including *Otx2* and *Lef1*. Consequently, TCF7L1 promotes pluripotency exit and suppresses epiblast lineage formation, thereby driving cells into PE specification. Conversely, deletion of *Tcf7l1* abrogates PE differentiation without restraining epiblast priming. Taken together, our study underscores the importance of transcriptional Wnt inhibition in regulating lineage segregation in ESCs and preimplantation embryo development as well as identifies TCF7L1 as key regulator of this process.

## Introduction

In early preimplantation development, the inner cell mass (ICM) cells face a binary decision between two distinct cell lineages: the naive pluripotent epiblast (EPI), which will form the embryo proper, and the extra-embryonic primitive endoderm (PE) cells, which will give rise to the endodermal component of the visceral and parietal yolk sac. On mouse embryonic day (E) 4.5, EPI and PE lineages are fully segregated, forming the expanded blastocyst, which is then ready for implantation in the uterus ^1–3^. Genetic studies have shown that the established EPI cell fate is centered on the transcriptional network mastered by NANOG, while PE fate is governed by GATA6 expression ^4–6^.

mESCs are the *in vitro* counterpart of the pluripotent preimplantation EPI and can be propagated indefinitely in pluripotent culture conditions ^7^. However, upon removal of self-renewing conditions, mESCs differentiate into post-implantation EPI-primed cells by first transiting through a formative pluripotent state ^8,9^. During this process, naive pluripotency transcription factors (TFs) such as *Nanog* and *Prdm14* are downregulated while formative pluripotency regulators such as *Lef1* and *Otx2* are induced together with the *de novo* methyltransferases *Dnmt3a* and *Dnmt3b* ^10,11^. Although mESCs resemble the preimplantation EPI, they can also spontaneously transit towards an extraembryonic PE-like state when cultured in naive pluripotency conditions ^12,13^, or by overexpression of the PE-specific genes *Gata6, Gata4 or Sox17* ^14–16^. The capacity of mESCs to commit into both EPI-primed and PE lineages provides a useful model to study the molecular mechanisms governing these early cell fate decisions ^12,13,17–19^.

Selective inhibition of the FGF/ERK pathway together with activation of the Wnt canonical pathway in mESC culture, using so-called 2i medium, promotes ground-state pluripotency ^20,21^, in which the PE-like population is absent ^13,20,22^. FGF/MAPK signaling has been proposed to be the key regulator of cell identity within the ICM ^4^. However, *Fgf4^-/-^* embryos express *Gata6*, which cannot be maintained after E3.25. This suggests that the onset of the PE program is FGF4-independent ^23–25^. Moreover, addition of FGF ligands, including FGF1 or FGF2, to mESC culture does not drive PE specification ^4, 26^, suggesting that alternative pathways may be implicated in this process.

Although several Wnt ligands and components are expressed in the preimplantation blastocyst ^27–33^, a specific role of the Wnt pathway in mammalian preimplantation development has not yet been described.Wnt signaling includes canonical or Wnt/β-catenin dependent and non-canonical or β-catenin-independent pathways ^34,35^. Repression of Wnt ligand secretion in naive mESCs, which simultaneously inhibits both canonical and non-canonical Wnt cascades, induces EPI-priming ^32^, yet interestingly, does not impact early lineage commitment at preimplantation stage ^10^. Exogenous activation of the Wnt/β-Catenin pathway is required for promoting mESC ground-state of self-renewal ^36,37^ indicating that pluripotent cells are receptive to external signaling modulation. However, the precise role of exogenous Wnt signaling inhibition in mESC lineage specification and in lineage segregation during preimplantation development is still unknown.

Upon Wnt ligand binding, unphosphorylated (active) β-Catenin accumulates in the cytoplasm. Subsequently, β-Catenin translocates to the nucleus, where it interacts with the T-cell factor/lymphoid enhancer factor (TCF/LEF) family of TFs to facilitate gene transcription. There are four TCF/LEF (TCF7, LEF1, TCF7L1 and TCF7L2) members in mammals. In mESCs, TCF7L1 is described as a transcriptional repressor, which limits the steady-state levels of the pluripotency network (*Oct4/Sox2/Nanog*) and of their common target genes ^38–40^. Alleviation of TCF7L1-mediated repression, following β-Catenin stabilization, is essential for pluripotency maintenance and self-renewal of mESCs ^41–43^. Interestingly, *Tcf7l1* deletion does not prevent the generation of primed epiblast stem cells (EpiSCs), even though it delays it ^8,40,44^, suggesting a minor role for TCF7L1 in EPI-priming. The role of TCF7L1 in commitment to other lineages remains unknown.

Here, we provide evidence that inhibition of Wnt signaling enhances PE cell fate commitment during preimplantation development as well as in mESC cultures. We demonstrate that forced expression of *Tcf7l1* efficiently drives mESCs towards a PE cell fate. By contrast, deletion of *Tcf7l1* in preimplantation embryos or in naive mESCs, abolishes their ability to differentiate into PE, without compromising epiblast priming. These TCF7L1-dependent effects are mediated by TCF7L1 binding and repression of essential genes for safeguarding naive pluripotency, such as *Nanog*, and *Prdm14*, causing exit from pluripotency. Notably, TCF7L1 also represses genes crucial for formative and EPI-primed programs including *Otx2, Lef1* and *Dnmt3b*, thereby preventing EPI priming and, hence, driving mESCs towards PE. These findings on the function of TCF7L1 during early development enhance our knowledge of the role played by the Wnt signaling pathway, and its key transcriptional players, the TCF/LEF factors, in regulating early developmental processes and provides further insights in the process of EPI vs. PE lineage segregation.

## Results

### EPI and PE lineages show differential Wnt/β-catenin pathway activities

Metastable subpopulations of naive EPI and PE-like cells arise in heterogenous mESC culture ^13,45^. The naive EPI and PE-like populations can be distinguished using platelet endothelial cell adhesion molecule 1 (PECAM1) and platelet-derived growth factor receptor α (PDGFRα) cell surface markers, respectively ^13^ (Fig. 1a). To identify the growth factors and signaling pathways involved in the regulation of EPI and PE lineage commitment in mESCs we analyzed publicly available RNA-sequencing (RNA-seq) data and evaluated expression differences in genes associated with naive pluripotency and PE between PDGFRα^-^/PECAM1^+^ (naive EPI) and PDGFRα^+^/PECAM1^-^ (PE-like) cells ^13^. As expected, naive EPI and PE-like cells expressed pluripotency- and PE-specific markers respectively (Fig. 1b). Gene ontology (GO) analysis of the differentially expressed genes (DEGs) between naive EPI and PE-like cells ^13^ highlighted differences in the regulation of the MAPK cascade and apoptosis pathways, in line with previous publications ^5,46–48^. Interestingly, GO analysis suggested a potential role of the canonical Wnt pathway in the regulation of EPI/PE lineage segregation (Fig. 1c; Supplementary Table 1). Compared with the PE-like population, naive EPI cells are characterized by a Wnt-ON state, defined by higher expression of well described positive Wnt regulators (*Fn1, Wnt7b, Lef1, Rspo1* and *Lgr4*) and lower levels of negative Wnt regulatory genes (*Gsk3b* and *Dkk1*) (Fig. 1d). Sorted PE-like population showed lower levels of total and active β-catenin (Fig. 1e), and lower levels of the transcriptional activators TCF7 and LEF1 (Supplementary Figure 1a) compared to naive EPI cells, supporting the notion that PE-like cells are characterized by limited Wnt signaling activity.

**Fig. 1.**
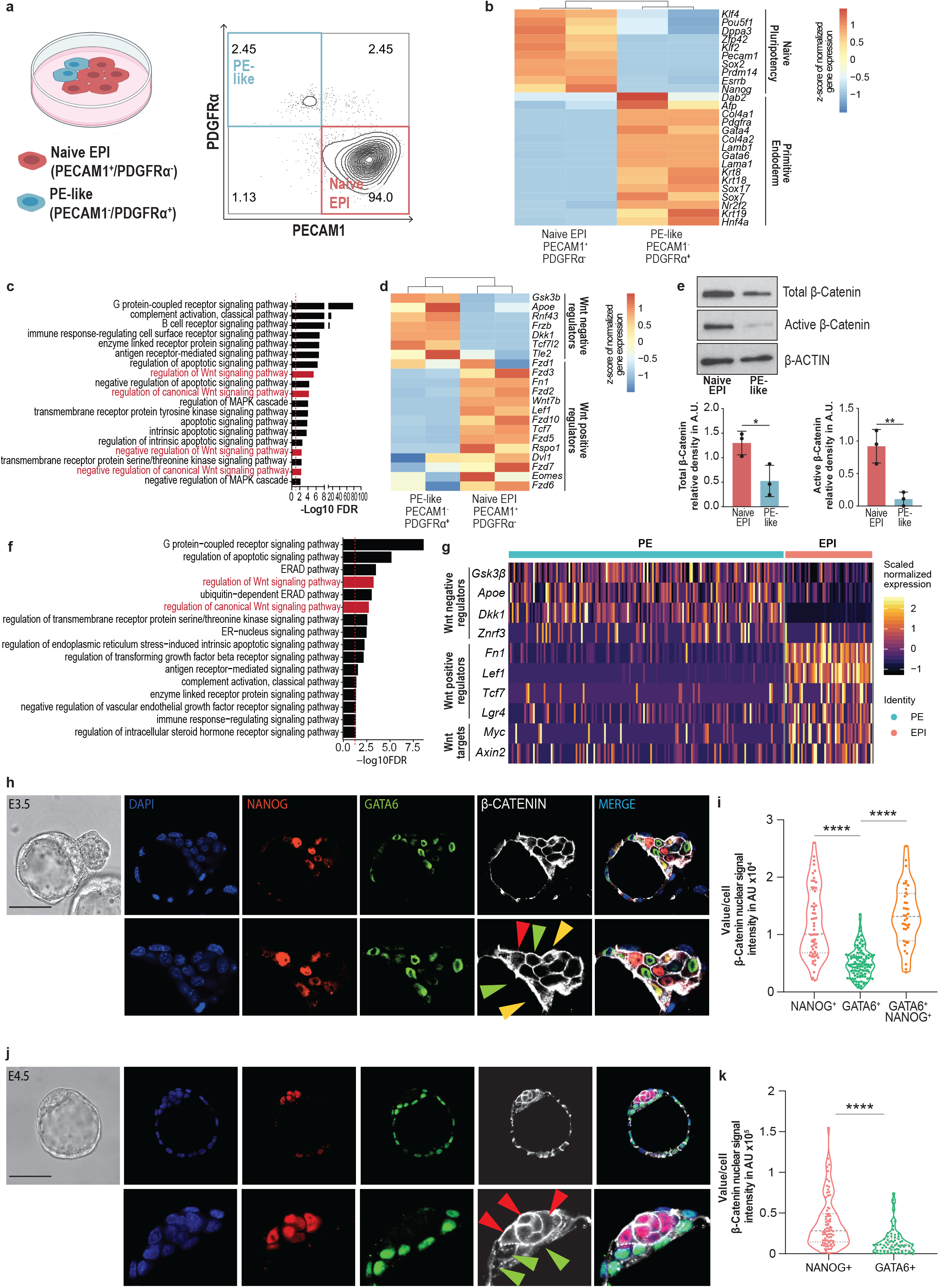
EPI and PE lineages correlate with opposite β-Catenin levels. (a) FACS density plot of mESC populations co-stained for PECAM1 and PDGFRα. (b) Expression of EPI and PE specific markers by respectively PECAM1^+^ and PDGFRα^+^ sorted cell populations. Unsupervised clustering. n=2 per condition ^13^. (c) GO enrichment analysis of DEGs between naive EPI and PE-like sorted cell populations ^13^. Red dashed line = FDR 0.05. (d) Expression of positive and negative Wnt regulators in naive EPI and PE-like sorted cell populations. Unsupervised clustering. n=2 per condition ^13^. (e) Representative images (up) and quantification of total/active β-catenin levels (down) in naive EPI and PE-like sorted cell populations. Mean ± SD; n=3; unpaired *t* test * p < 0.05, ** p < 0.01. (f) GO enrichment analysis of DEGs between EPI and PE cells in E4.5 embryos from ^51^. Red dashed line = FDR 0.05. (g) Expression of positive and negative Wnt regulators in EPI and PE lineages in E4.5 embryos ^51^. (h) Representative immunofluorescence (IF) image of active β-catenin signal in E3.5 embryo. EPI-NANOG^+^ cells (red arrow), PE-GATA6^+^ cells (green arrow) and double-positive NANOG^+^/GATA6^+^ cells (yellow arrow). Scale bar=50μM. Zoomed region of interest is reported below the images. (i) Nuclear active β-catenin signal in EPI-NANOG^+^, PE-GATA6^+^ and doublepositive NANOG^+^/GATA6^+^ cells. Integrated intensity in arbitrary units (AU). Mean ± SEM; NANOG^+^ n=38, GATA6^+^ n=111 and NANOG^+^/GATA6^+^ cells n=58; 2 independent experiments. One-way ANOVA, *p<0.05, ***p<0.001. (j) Representative IF image of active β-catenin signal in freshly isolated E4.5 embryo. EPI-NANOG+ cells (red arrow), PE-GATA6+ cells (green arrow). Scale bar=50μM. Zoomed region of interest is reported below the images. (k) Nuclear active β-catenin signal in EPI-NANOG+ and PE-GATA6+ cells. Integrated intensity in arbitrary units (AU). Mean ± SD; NANOG+ n=78; GATA6+ n=68; 2 independent experiments; t test ****p<0.0001.

Previous studies reported that the canonical Wnt pathway is transcriptionally active in the ICM of preimplantation embryos by using an *Axin2* transcriptional reporter ^32^ or by immunofluorescence for active β-Catenin ^30,49^. However, this did not address whether EPI and PE lineages display differential activity levels of canonical Wnt pathway. Using publicly available single cell RNA-seq (scRNA-seq) analysis of two independent E4.5 preimplantation embryo data sets ^50,51^, we confirmed the differential activity of the Wnt/β-Catenin pathway in EPI and PE lineages (Fig. 1f, Supplementary Fig. 1b-d, and Supplementary Table 2, 3). EPI cells expressed higher levels of canonical Wnt positive regulators (*Fn1, Lef1* and *Lgr4*) and Wnt targets (*Axin2* and *Myc*) compared to PE cells. By contrast, PE cells expressed higher levels of genes involved in Wnt pathway repression (*Gsk3b, Znfr3* and *Dkk1*) (Fig. 1g and Supplementary Fig. 1e). Therefore, we explored the levels of active β-Catenin in EPI (NANOG^+^) and PE (GATA6^+^) cells in preimplantation embryos at E3.5 and E4.5. Three distinct subpopulations of single-positive NANOG^+^ (EPI), single-positive GATA6^+^ (PE) and doublepositive NANOG^+^/GATA6^+^ cells were identified in the ICM of freshly isolated E3.5 blastocysts (Fig. 1h, i). By E4.5, the ICM consisted of only two distinct populations: single-positive NANOG^+^and single-positive GATA6^+^ cells in both E4.5 freshly isolated and E3.5+24H *ex vivo* cultured embryos (Fig. 1j and Supplementary Fig. 1f). Although active β-Catenin was primarily localized at the cell membrane, in agreement with its function in adherent junctions ^52^, we also detected diffused cytoplasmatic and nuclear staining for active β-Catenin (Fig. 1h, j and Supplementary Fig.1f), in accordance with previous reports ^30^. Quantification of nuclear and total (membrane+intracellular) active β-Catenin confirmed its lower levels in GATA6^+^ cells (green arrows) compared to NANOG^+^ cells (red arrows) and co-expressing NANOG^+^/GATA6^+^ cells (yellow arrows) at E3.5 or E4.5 blastocyst stages (Fig. 1i, k and Supplementary Fig. 1f-h).

### Wnt signaling inhibition promotes formation of the PE lineage

Previous studies demonstrated an important role of Wnt signaling during post-implantation development or mESC self-renewal ^20,21,32,53–57^. Although blocking Wnt ligand secretion in mESC promotes EPI-priming ^58^, the effect of paracrine canonical Wnt inhibition on mouse early lineage specification has not been explored. We therefore assessed if modulation of the canonical Wnt signaling pathway would alter the proportion of naive EPI and PE-like cells *in vitro* as well as within the ICM of mouse embryos. Treatment of mESCs with the physiological canonical Wnt antagonist Dickkopf1 (DKK1) decreased active β-Catenin protein levels (Supplementary Fig. 2a), causing a significant enrichment in the percentage of PDGFRα^+^/PECAM1^-^ (PE-like) cells (Fig. 2a), along with a notable upregulation of PE gene markers, including *Gata6, Gata4* and *Sox17* (Fig. 2b). This suggested a direct role of Wnt signaling inhibition in PE formation. We also cultured E2.5 embryos for 48 hours to the late blastocyst stage in the presence or absence of DKK1 (E2.5+48H). Lineage assessment showed that DKK1-treated embryos contained an increased number of PE (GATA6^+^) cells, and a decreased number of EPI (NANOG^+^) cells, compared to controls (Fig. 2c, d). While control embryos showed a proportion of around 50% of NANOG^+^ or GATA6^+^ cells inside the ICM at E2.5+48H, DKK1-treated embryos displayed 70% of GATA6^+^ cells (Fig. 2e), demonstrating preferential PE commitment upon Wnt inhibition. Interestingly, DKK1-treated embryos showed an accelerated development with more hatched blastocysts at E2.5+48H (Supplementary Fig. 2b). A direct correlation between PE specification and blastocyst lumen expansion has been recently shown ^59^. In agreement, the blastocyst lumen volume of DKK1-treated embryos was 72% larger (Fig. 2f, g), which was associated with a significant increase of 80% in the embryo size area compared to control embryos (Fig. 2h, i and Supplementary Fig. 2c). Thus, these results demonstrate a direct link between Wnt pathway inhibition and PE commitment at the preimplantation stage.

**Fig. 2.**
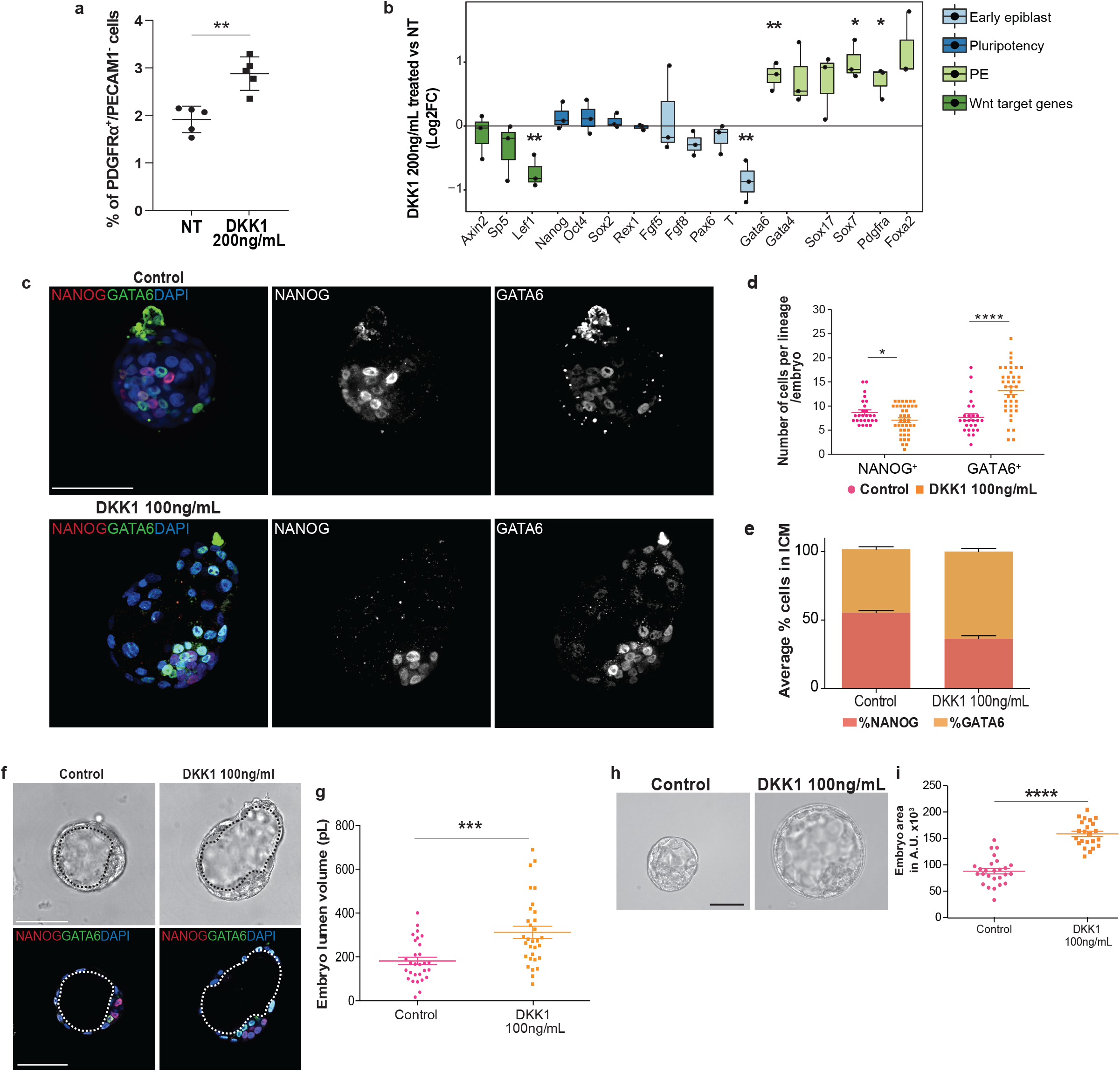
Wnt signaling inhibition promotes formation of the PE lineage. (a) Percentage of PE-like cells (PDGFRα^+^/PECAM1^-^) in control and DKK1-treated mESCs. Mean ± SD; n=5, t test **p<0.01. (b) Gene expression analysis of early EPI, pluripotency, PE and Wnt target genes on control and DKK1-treated mESCs. Mean ± SD; n=3; *t* test *p<0.05, **p<0.01. (c) Representative IF image of NANOG and GATA6 protein signals in E2.5+ 48H control and DKK1 treated embryo. Nuclei were counterstained with DAPI. Scale bar = 50μM. (d) Number of NANOG+ and GATA6+ cells (counts) per embryo. Mean ± SEM; Control: 26; DKK1 n=39; 3 independent experiments. *t* test *p<0.05, ****p<0.0001. (e) Percentage of NANOG^+^ and GATA6^+^ cells normalized on total number of ICM per embryo. Mean ± SEM; Ctrl n=28, DKK1 n=42; 3 independent experiments. (f) Representative BF and IF image of E2.5+ 48H control and DKK1 treated embryo. Nuclei were counterstained with DAPI. Black and white dotted lines delimitate blastocyst cavity (lumen). Scale bar=50 μM. (g) Embryo lumen volume reported in pico liters (pL). Mean ± SEM; Ctrl n=29, DKK1 n=31; 3 independent experiments. *t* test ***p<0.001. (h) Representative brightfield (BF) image of embryo morphology. Scale bar = 50μM. (i) Embryo total area reported in A.U. Mean ± SEM; Ctrl n=26, DKK1 n=23; 3 independent experiments. *t* test ****p<0.0001.

### *Tcf7l1* deletion impairs EPI to PE transition without permanently compromising pluripotency exit and commitment to neuroectodermal lineage

To investigate whether PE specification *in vitro* is mediated by TCF/LEF transcriptional activity, we used the inhibitor of β-catenin–responsive transcription iCRT3, which prevents the interaction of β-catenin and TCF factors, thus inhibiting transcription of Wnt target genes ^60^. iCRT3-treated mESCs exhibited a significantly higher percentage of PDGFRα^+^/PECAM1^-^ (PE-like) cells (Fig. 3a). We observed a concomitant enhanced expression of PE markers compared to control (Fig. 3b), indicating that repression of TCF/LEF transcriptional activity is directly involved in EPI to PE specification.

**Fig. 3.**
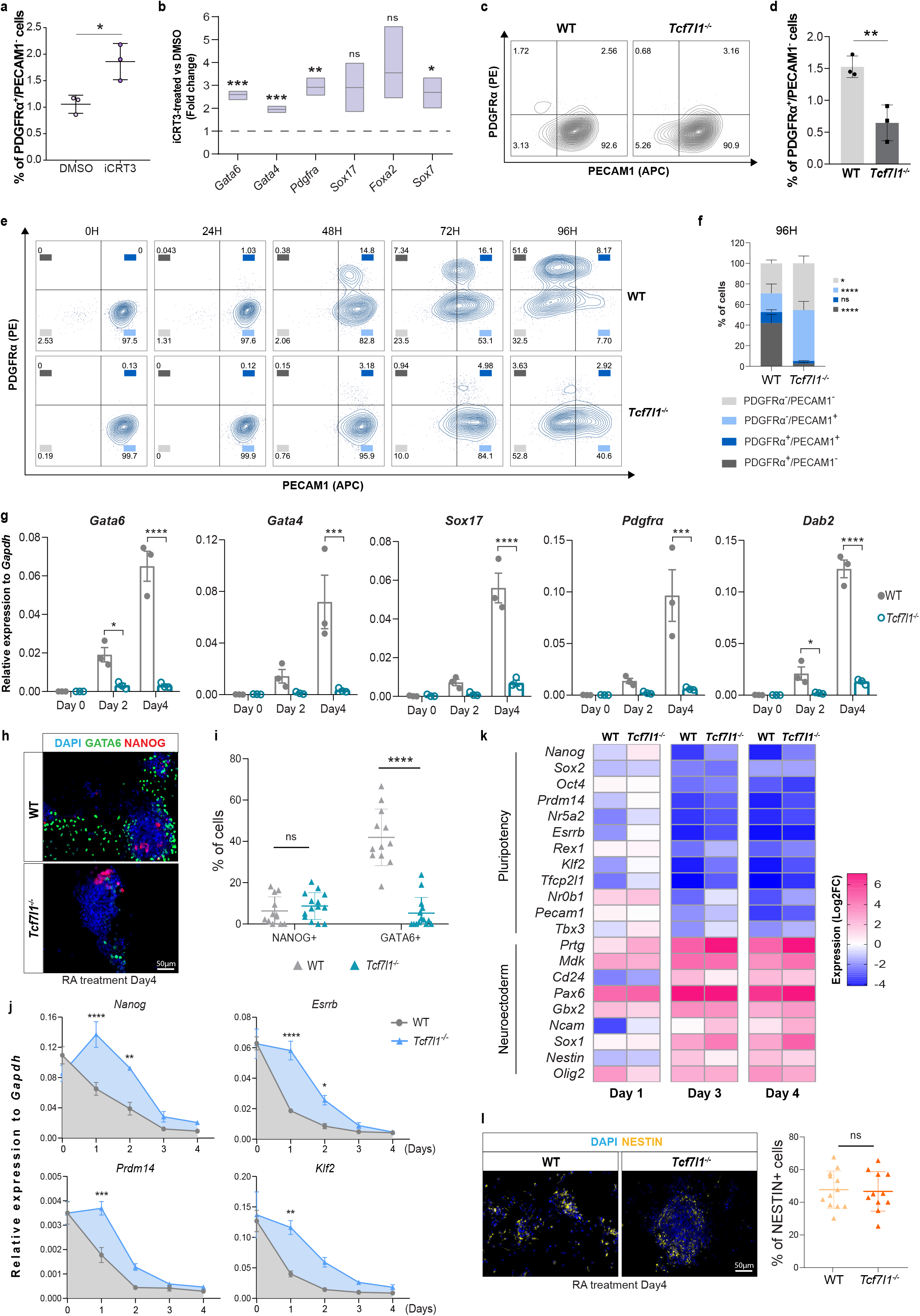
*Tcf7l1* deletion specifically impairs EPI to PE transition without compromising commitment to embryonic lineages. (a) Percentage of PDGFRα^+^/PECAM1^-^ cells revealed by flow cytometry analysis in mESC treated for 3 passages with 10μM iCRT3. Mean ± SD; n=3; *t* test p*< 0.05. (b) qRT-PCR of extraembryonic endoderm markers in mESCs cultured with 10μM iCRT3. Values represent expression fold change relative to DMSO (control) samples. Fold Change ± SD; n=3; multiple *t* test *p<0.05, **p<0.01, p***<0.001. (c) Representative flow cytometry images of mESC populations co-stained for PECAM1 and PDGFRα in WT and *Tcf7l1^-/-^* cells. (d) Flow cytometry quantification of PDGFRα^+^/PECAM1^-^ cells in WT and *Tcf7l1^-/-^* mESCs. Mean ± SD; n=3; *t* test **p<0.01. (e) Representative flow cytometric images of mESC populations co-stained for PECAM1 and PDGFRα markers in WT and *Tcf7l1^-/-^* cells upon 0, 24, 48, 72 and 96H of 0.25μM RA treatment. n=3. (f) Flow cytometry analysis of PDGFRα and PECAM1 populations at 96H of RA treatment. Mean ± SD; n=3; two-way ANOVA test *p<0.05, **** p<0.0001. (g) qRT-PCR showing relative to *Gapdh* gene expression of extraembryonic markers in WT and *Tcf7l1^-/-^* mESCs upon 0, 2 and 4 days of RA treatment. Mean ± SEM; n=3; two-way ANOVA test *p<0.05, ***p<0.001, **** p<0.0001. (h) Representative IF image of GATA6 and NANOG in WT and *Tcf7l1^-/-^* mESCs upon 4 days of RA. Scale bar=50μm. (i) Quantification of NANOG+ and GATA+ cells of Figure 3H. Mean ± SD; Mann-Whitney test ****p<0.0001. Each dot represents % of cells in a field of view; WT: 12, *Tcf7l1^-/-^* : 14. (j) Gene expression analysis of pluripotency markers in WT and *Tcf7l1^-/-^* mESCs upon 0, 24, 48, 72 and 96H of 0.25Mm RA treatment. Mean ± SEM; n=3; two-way ANOVA test **p<0.01, ***p<0.001, **** p<0.0001. (k) Representative IF image of NESTIN in WT and *Tcf7l1^-/-^* mESCs upon 4 days of RA and quantification of NESTIN+ cells. Mean ± SD; Mann-Whitney test, ns= no significant differences. Each dot represents % of cells in a field of view; WT: 12, *Tcf7l1^-/-^* : 11. (l) Gene expression analysis of embryonic neuroectodermal markers in WT and *Tcf7l1^-/-^* mESCs upon 1, 3 and 4 days of RA treatment. Gene expression values are reported as Log2 of fold change expression. n=3.

TCF7 and TCF7L1 are the most abundant TCF/LEF members in mESCs ^61,62^. While TCF7 is considered a Wnt transcriptional activator ^63^, TCF7L1 is primarily a transcriptional repressor ^40,64^. To assess their role in PE formation, we evaluated the effects of *Tcf7* or *Tcf7l1* deletion on mESC heterogeneity. While deletion of *Tcf7l1* resulted in a significant reduction of the PDGFRα^+^/PECAM1^-^ cell population (Fig. 3c and 3d), the fraction of PE-like cells was not affected by deletion of *Tcf7* (Supplementary Fig. 3a and 3b).

To further explore the role of TCF7L1 and TCF7 in PE differentiation, WT, *Tcf7l1^-/-^* and *Tcf7^-/-^* mESCs were firstly adapted in 2i medium plus LIF in which the transcriptome of ESCs closely resembles that of ICM cells ^6^. Next, we differentiated the cells using Retinoic Acid (RA) in basal medium, which is known to promote pluripotency exit and induce both extraembryonic PE and embryonic neuroectodermal differentiation ^65^. WT cells treated with RA progressively transited towards the PE fate (PDGFRα^+^/PECAM1^-^) yielding more than 50% PE cells by 4 days (Fig. 3e). Strikingly, although *Tcf7l1^-/-^* cells cultured with RA lost expression of the PECAM1 pluripotency marker, they did not differentiate into PE-like cells (Fig. 3e, f) and PE marker gene expression was not induced in RA-treated *Tcf7l1^-/-^* cells (Fig. 3g). Furthermore, the percentage of GATA6^+^ cells was considerably lower in *Tcf7l1^-/-^ cells* treated with RA (Fig. 3h, i). On the contrary, *Tcf7*^-/-^ cells were able to undergo PE specification after RA treatment, as shown by the high expression of PE markers, which was accompanied by pluripotency exit (Supplementary Fig. 3c-e).

TCF7L1 binds and represses pluripotency markers, suggesting that the *Tcf7l1*-dependent PE differentiation defect might be caused by impaired pluripotency exit ^40,63^. Although decrease of pluripotency gene expression was delayed in *Tcf7l1^-/-^* cells at early differentiation time points (D1 and D2) as previously reported ^8^, *Tcf7l1^-/-^* cells were able to efficiently exit pluripotency after D3 of RA treatment similarly to WT (Fig. 3i-k). To evaluate whether *Tcf7l1*-dependent differentiation impairment was related to other lineages we investigated the expression of neuroectodermal markers which is also induced by RA treatment. Interestingly, both WT and *Tcf7l1^-/-^* mESCs exhibited strong and similar upregulation of neuroectodermal genes and showed equal NESTIN protein levels ^66^ (Fig. 3k, l).

In summary, we demonstrated that *Tcf7l1* deletion in mESCs delays but does not stall pluripotency exit. Furthermore, *Tcf7l1^-/-^* cells increase neuroectodermal marker expression to the same level as WT cells, suggesting that *Tcf7l1^-/-^* cells maintain their embryonic differentiation potential. In contrast, *Tcf7l1*^-/-^ mESCs are incapable of converting to PE-like cells, illustrating an essential role of TCF7L1 in extraembryonic PE induction *in vitro*.

### *Tcf7l1* expression is sufficient to drive PE cell fate in ESCs

Forced expression of PE-lineage specific TFs ^14–16^ or repression of naive EPI-specific TFs ^67,68^ promotes mESC differentiation towards PE. However, the putative role of non lineage-specific TFs in PE formation has yet to be studied. To examine the role of TCF7L1 in PE cell fate specification in mESCs, we performed a gain-of-function analysis using a doxycycline (Dox)-inducible (Tet-OFF) *Tcf7l1* ESC line ^69^. Following Dox removal for 24H, we observed robust expression of the TCF7L1-FLAG tag protein along with Venus YFP (Supplementary Fig. 4a, b). However, not all cells responded equally to the Dox removal as it can be seen from YFP fluorescence intensity variations (Supplementary Fig. 4b), possibly due to time and cellular variations of response to Dox removal ^70^. We confirmed that *Tcf7l1* overexpression correlated with Venus^high^ expression (Supplementary Fig. 4c). To study the effect of uniform overexpression (OE) of *Tcf7l* in mESC culture, we sorted Venus^high^ (Supplementary Fig. 4c) cells 24H after Dox removal (D1) and replated the cells for an additional 7 days under ESC self-renewal conditions. By day 4, Venus^high^ cells exhibited a flat morphology reaching an epithelial-like morphology by day 8 (Fig. 4a). After 48H of *Tcf7l1* induction, we observed progressive upregulation of the PE genes *Gata6, Gata4, Pdgfrα* and *Sox17*, reaching levels comparable to those of embryo-derived extraembryonic endoderm stem (XEN) cells by day 8 (Fig. 4b) along with a decreased expression of the pluripotency genes *Nanog* and *Sox2* (Fig. 4c). Furthermore, after 6 days of *Tcf7l1* induction, mESCs exhibited increased GATA6 protein levels, which was accompanied with decreased NANOG levels (Fig. 4d,e and Supplementary Fig. 4d). By contrast, forced expression of *Tcf7* (Supplementary Fig. 4e, f), resulted in negligible changes in PE gene expression (Supplementary Fig. 4g), indicating that TCF7 has a minor role, if any, in PE lineage commitment.

**Fig. 4.**
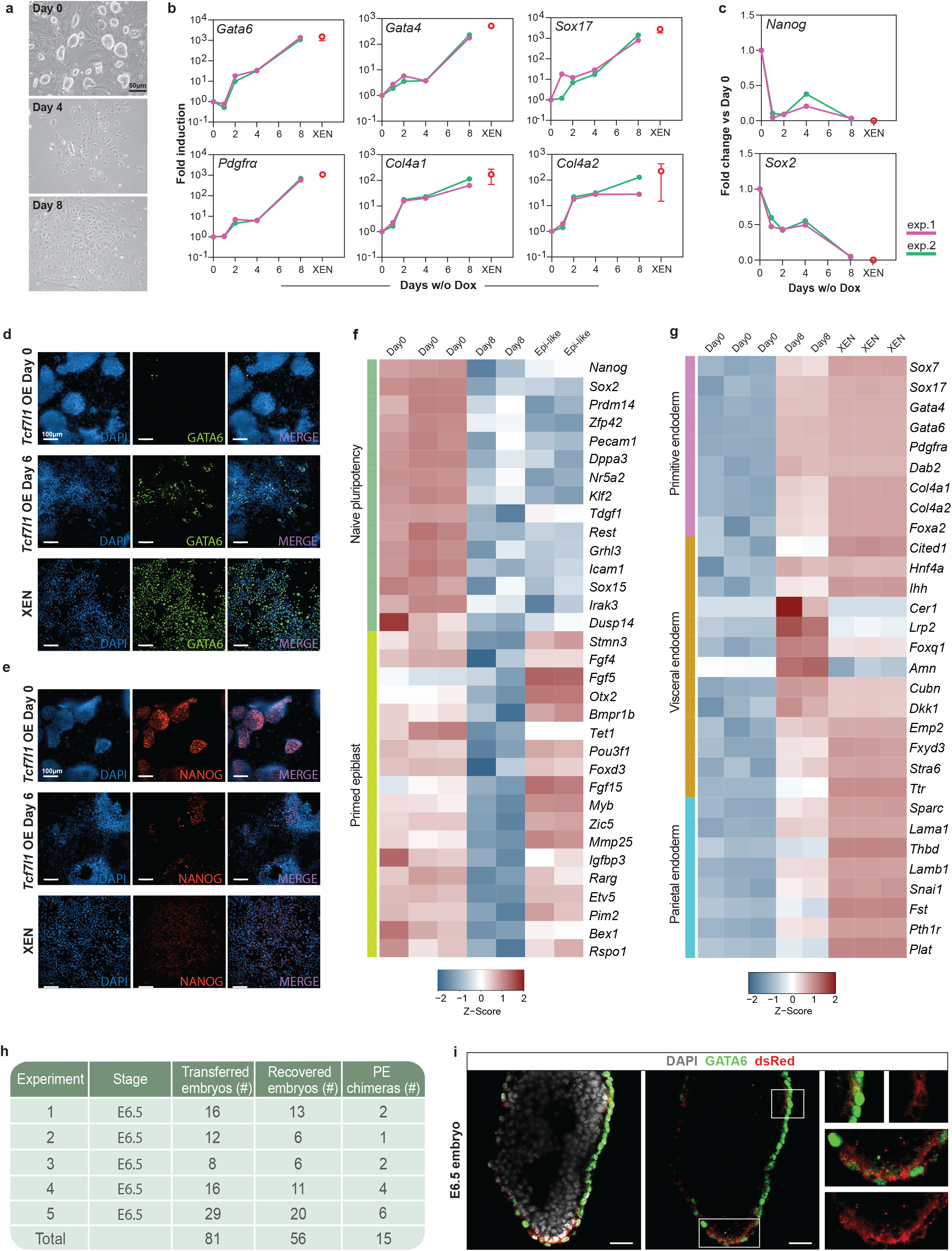
*Tcf7l1* overexpression promotes PE commitment of ESCs. (a) Representative BF images of *Tcf7l1*-OE-mESCs cultured with Dox (D0) or without Dox (D4 and D8). Scale bar = 50μm. (b) Gene expression analysis of PE markers during *Tcf7l1* OE in mESCs. Two independent experiments (pink and green) are shown at indicated time points relative to uninduced mESCs alongside XEN cells (red circle). For XEN, mean ± SD; n=2. (c) Gene expression analysis of pluripotency markers during *Tcf7l1* overexpression in mESCs as in Figure 4b. For XEN, mean ± SD; n=2. (d) Representative IF image of GATA6 in *Tcf7l1*-OE-mESCs after 0 and 6 days of induction and XEN cells. Scale bar = 100μM. (e) Representative IF of NANOG as in Figure 3d. (f) Normalized gene counts heatmap showing expression of naive pluripotency and primed epiblast markers in mESCs (D0), after 8 days of *Tcf7l1* induction and in Epi-like cells. (g) Normalized gene counts heatmap showing expression of primitive endoderm, visceral and parietal endoderm markers in mESCs (D0), after 8 days of *Tcf7l1* induction and XEN cells. (h) Chimeric contribution of Venus^high^/dsRed^+^ to PE-derived lineages. (i) Chimeric E6.5 embryo stained with GATA6 and dsRed. Scale bar=50μM. Zoomed regions of interest are reported adjacent to the images.

Next, we performed RNA-seq of D0 and D8 *Tcf7l1*-OE cells to assess gene expression patterns resulting from *Tcf7l1* gain-of-function. We identified 2063 DEGs between D0 and D8 (|log2FC|>2, FDR≤0.05) (Supplementary Table 4). GO analysis of DEGs showed association with embryo morphogenesis and endoderm differentiation processes (Supplementary Fig. 4h and Supplementary Table 4). In addition, gene expression profile of D8 *Tcf7l1*-OE cells showed that, unlike Wnt secretion inhibition which promotes EPI-priming ^32^, overexpression of *Tcf7l1* reduced primed and naive pluripotency related gene expression (Fig. 4f). Visceral endoderm (VE) and Parietal endoderm (ParE) are major derivatives of primitive endoderm and are distinguished by characterized markers ^71–75^. Interestingly, while XEN cells transcriptionally resemble more ParE ^72,73,76^, *Tcf7l1*-OE-mESCs displayed high levels of PE and VE markers expression suggesting that OE of *Tcf7l1* promotes a VE-rather than a ParE-cell commitment (Fig. 4g).

Upon blastocyst injection, mESC-derived PE cells contribute to the extraembryonic layers of mouse post-implantation embryos ^13,73^. Therefore, to test whether *Tcf7l1*-OE-mESCs can contribute to PE, we injected them into preimplantation blastocysts. To trace the descendants during development, cells were transfected with a PiggyBac-based vector encoding for a constitutive dsRed protein (PB-dsRed)^77^. Sorted Venus^high^/dsRed^+^ cells were harvested 4 days after Dox-removal and injected into E3.5 WT host blastocysts, which were then transferred into pseudopregnant recipient mice. After E6.5 embryos were isolated, dsRed^+^ cells were clearly visible in chimeric embryos (Fig. 4h and Supplementary Fig. 4i). Whole embryo immunofluorescence co-staining for dsRed and GATA6, showed co-expression of the VE marker GATA6 ^75^ in dsRed+ cells supporting the VE fate of the descendants of the injected cells (Fig. 4i and Supplementary Fig. 4j). Overall, these findings demonstrate that upon forced expression of *Tcf7l1*, mESCs engage in PE gene activation, creating PE-like cells that can contribute to the VE lineage *in vivo*.

### Differentiation to PE fate upon *Tcf7l1* overexpression follows downregulation of naive and formative pluripotency programs

To define the transcriptional dynamics regulating early embryo development and PE formation, we performed transcriptomic analysis of *Tcf7l1*-OE-mESCs at different timepoints. *Tcf7l1*-OE-mESCs were collected 24H, 48H and 96H after Dox removal along with Dox-treated cells as control for RNA-seq analysis. Gene expression profiling identified 856 DEGs on D1, 1663 DEGs on D2 and 1656 DEGs on D4 between *Tcf7l1*-OE and Dox-treated mESCs (Supplementary Table 5). GO analysis of downregulated genes after 1 day of *Tcf7l1*-OE showed enrichment of pathways involved in the regulation of embryonic development, cell fate commitment and negative regulation of cell adhesion along with the known developmental pathways, BMP and FGF (Fig. 5a). In addition, GO terms and KEGG pathways related to stem cell development and differentiation together with terms associated with canonical Wnt pathway were identified in D1 *Tcf7l1*-OE-mESCs (Fig. 5a and Supplementary Table 6). Specifically, Wnt target genes (*Axin2, Lef1 and Sp5*) as well as genes associated with naive and general pluripotency (*Nanog, Klf2, Prdm14, Dpp4 and Fn1*) were downregulated. In line with previous reports ^78^, we found little effect on *Oct4* expression on D1, demonstrating that PE precursors retain *Oct4* expression at early stages ^13,79^.

**Fig. 5.**
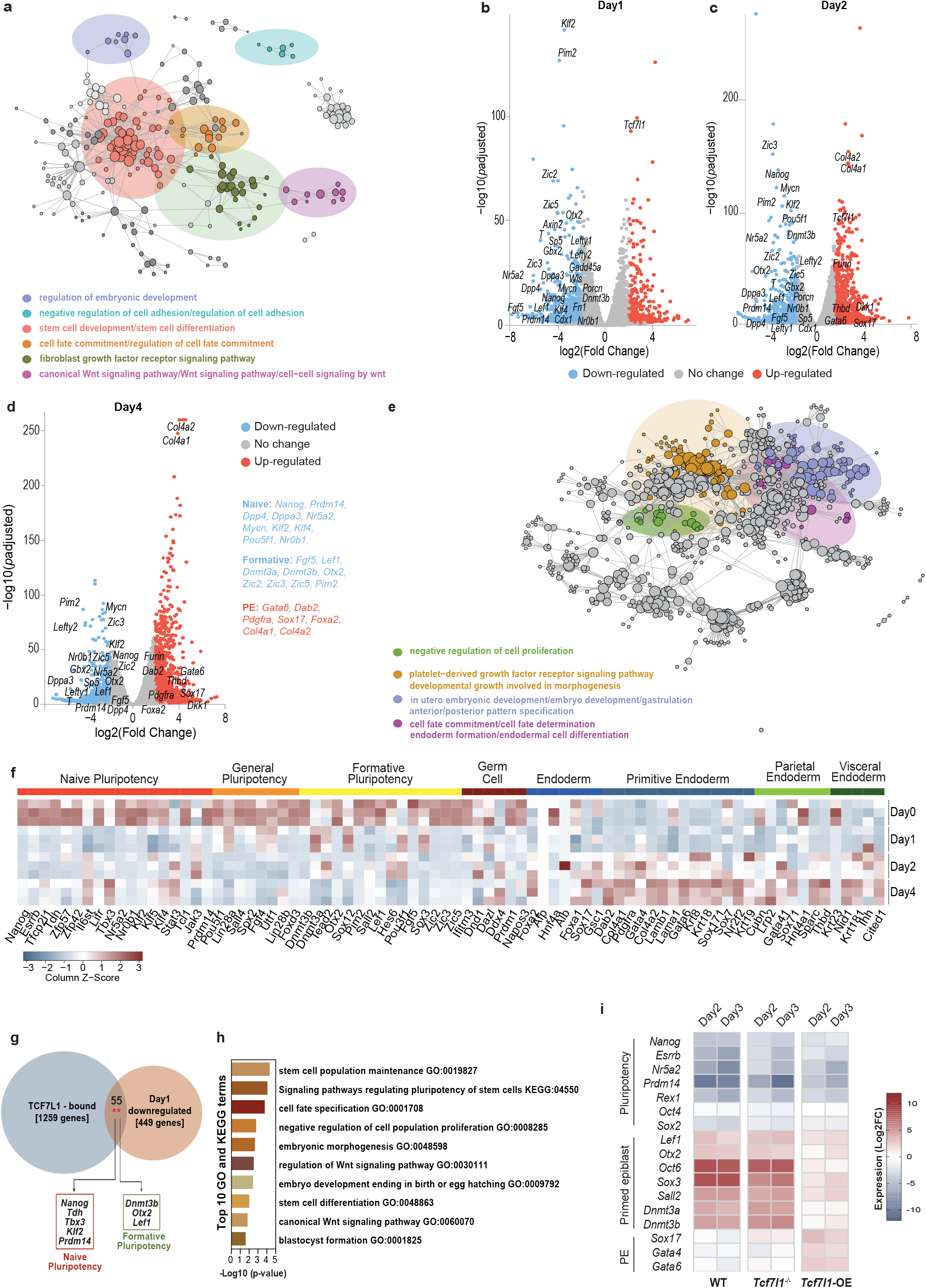
*Tcf7l1* overexpression promotes conversion to PE fate through a downregulation of naive and formative pluripotency. (a) GO and KEGG functional enrichment analysis for *Tcf7l1*-OE-mESCs after 24h hours of induction. Functional term networks were obtained using Cytoscape’s ClueGO plugin. Each node corresponds to an enriched term and edges connect terms that share a significant number of genes. Network communities (shown in color) were created using the Louvain algorithm, adjusted for betweenness centrality. (b) Volcano plot showing down- (blue dots) and upregulated (red dots) genes altered in *Tcf7l1*-OE-mESCs after 24h hours of induction. Gene values are reported as a Log2FoldChange (adjusted p-value<0.05). Annotated points correspond to selected marker genes. (c) Volcano plot showing DEGs in *Tcf7l1*-OE-mESCs after 2 days of induction as in Figure 5b. (d) Volcano plot showing DEGs in *Tcf7l1*-OE-mESCs after 4 days of induction as in Figure 5b. (e) GO and KEGG functional enrichment analysis for *Tcf7l1* mESCs after 4 days of induction. Network was created and analyzed as described in Figure 5a. (f) Normalized gene counts heatmap showing 80 selected specific lineage markers in WT and *Tcf7l1^-/-^* cells upon 1, 2 and 4 days of induction. (g) Venn diagram indicating genes bound by TCF7L1 (1259 genes) from publicly available ChIP-seq data ^37^ and downregulated genes in *Tcf7l1*-OE-mESCs after 1 day of induction (449 genes with unique promoters). There were 55 genes shared between the two sets (p-value of a Fisher’s exact test = 0.0021) which include markers of naive and formative pluripotency. (h) Top 10 functional enrichment terms for the 55 genes downregulated in *Tcf7l1*-OE-mESCs after 1 day of induction and bound by TCF7L1. (i) Gene expression analysis of pluripotency, primed epiblast/formative, and PE gene markers in WT, *Tcf7l1^-/-^* and *Tcf7l1*-OE-mESCs upon EpiLCs differentiation. Gene expression values are reported as Log2 of fold change expression.

Recently, formative pluripotency has been described as an intermediate phase that precedes primed epiblast formation upon naive pluripotency exit ^9,11,80^. The formative state is characterized by decreased expression levels of naive pluripotency markers and increased expression of peri-implantation markers such as *Lef1* and *Dmnt3a* and *Dnmt3b*. Expression of *Otx2* has also been shown to be essential for the maintenance of the formative pluripotency state ^80–82^. Overexpression of *Tcf7l1* caused a decrease in expression of several formative-specific genes including *Otx2, Lef1, Dnmt3b, Fgf5, Pim2 and Zic2* ^83^ already on D1, which was maintained on D2 and D4 (Fig. 5b-d).

We did not observe increased expression of endoderm genes on D1 (Fig. 5b). However, on D2, endodermal genes were induced, which was even more obvious 4 days after OE of *Tcf7l1* (Fig. 5c, d). Upregulated genes on D4 were associated with the formation of endoderm, cell fate determination, the PDGFRα signaling pathway as well as gastrulation and anterior/posterior pattern specification (Fig. 5e and Supplementary Table 7). We observed increased levels of primitive, parietal and visceral endoderm genes, including master regulators of endoderm formation (*Gata6, Gata4* and *Sox17*), genes involved in cell adhesion (*Col4a1, Col4a2 and Dab2*) and other genes associated with PE function (*Pdgfrα, Sparc, Thbd, Nid1* and *Cited1* (Fig. 5f). These findings demonstrate that activation of PE genes follows the downregulation of naive and formative pluripotency.

Using a TF-gene target enrichment analysis ^84,85^, we found that the genes downregulated after *Tcf7l1* OE were putative targets of TCF7L1, CTNNB1 (β-Catenin), NANOG, SOX2 and OCT4 (Supplementary Fig. 5a), confirming the coregulation of TCF7L1 targets with the pluripotency TF network ^86^.

To further assess whether TCF7L1 directly regulates formative pluripotency gene expression, we integrated RNA-seq data of genes downregulated on D1 in *Tcf7l1*-OE-mESCs with publicly available TCF7L1 ChIP-seq data on mESCs ^37^. This revealed that 55 genes (p=0.0021) that were downregulated on D1, were also bound by TCF7L1 around the TSS (transcription start site) region (Fig. 5g, Supplementary Fig. 5b and Supplementary Table 8). Common genes included naive (*Nanog, Tdh, Tbx3, Klf2 and Prdm14*), and formative (*Dnmt3b, Otx2, Sox12* and *Lef1*) pluripotency regulators. Interestingly, bound genes were associated with stem cell population maintenance, blastocyst formation and embryonic development together with signaling pathways regulating pluripotency and canonical Wnt signaling (Fig. 5h and Supplementary Table 9).

Hence, our results demonstrate direct repression of naive and formative pluripotency genes by TCF7L1, which might prevent formative- and consequently EPI-priming. To test this hypothesis, we performed RT-qPCR gene expression analysis of WT, *Tcf7l1^-/-^* and *Tcf7l1*-OE mESCs during epiblast-like cell (EpiLCs) differentiation ^9^. WT and *Tcf7l1^-/-^* mESCs successfully downregulated naive and general pluripotency markers and upregulated key genes involved in formative and primed EPI differentiation ^80^ (Fig. 5i). *Tcf7l1*-OE cells showed comparable downregulation of naive and general pluripotency genes, yet a decreased upregulation of formative genes compared to WT and *Tcf7l1^-/-^* cells (Fig. 5i). Interestingly, *Tcf7l1*-OE cells displayed increased expression of PE genes in EpiLCs differentiation conditions, indicating that forced expression of *Tcf7l1* has a dominant effect, promoting PE differentiation even in non-permissive PE cell culture conditions (Fig. 5i). These results confirm the role of TCF7L1 as a barrier to formative and epiblast differentiation and as the driving force to PE specification.

### TCF7L1 is required for PE cell fate specification during *in vivo* preimplantation development

Using publicly available scRNA-seq and single-cell gene regulatory inference and clustering (pySCENIC) data from mouse embryos ^87^, we assessed the activity of TCF/LEF TFs along with their target genes (known as regulons) at the preimplantation stage. Interestingly, regulon analysis predicted that Tcf7- and Lef1-regulons were transcriptionally active primarily in the EPI compartment, whereas Tcf7l1- and Tcf7l2-regulons appeared transcriptionally active in PE cells (Fig. 6a, b). To unveil the role of TCF/LEF factors during PE cell lineage commitment, we used published scRNA-seq data, where the Harmony algorithm combined with Palantir were used to construct a spatio-temporal map of PE specification in mouse embryos ^51^. *Tcf7l1* and *Tcf7l2* showed distinct expression patterns during preimplantation development. Specifically, we found a transient upregulation (pulse) of *Tcf7l1* in PE-fated cells between E3.5 and E4.5, which is compatible with involvement of TCF7L1 in PE formation (Fig. 6c). In line with our previous data on mESCs, naive and formative pluripotency markers were significantly decreased, whereas PE genes were considerably induced, following the *Tcf7l1* pulse (Fig. 6c). By contrast, EPI-fated cells lacked the *Tcf7l1* pulse in gene expression levels (Supplementary Fig. 6a). To further elucidate the role of TCF/LEF factors during development, E2.5 embryos were treated *ex vivo* with iCRT3 for 48H (Fig. 6d). This resulted in embryos with a significantly increased number of GATA6^+^ cells, fewer NANOG+ cells (Fig. 6e, f) and a larger lumen volume (Fig. 6g), phenocopying the effects exerted by DKK1 (Fig. 2). This suggests that TCF factors play an essential role in regulating developmental processes such as cavitation and PE cell fate specification.

**Fig. 6.**
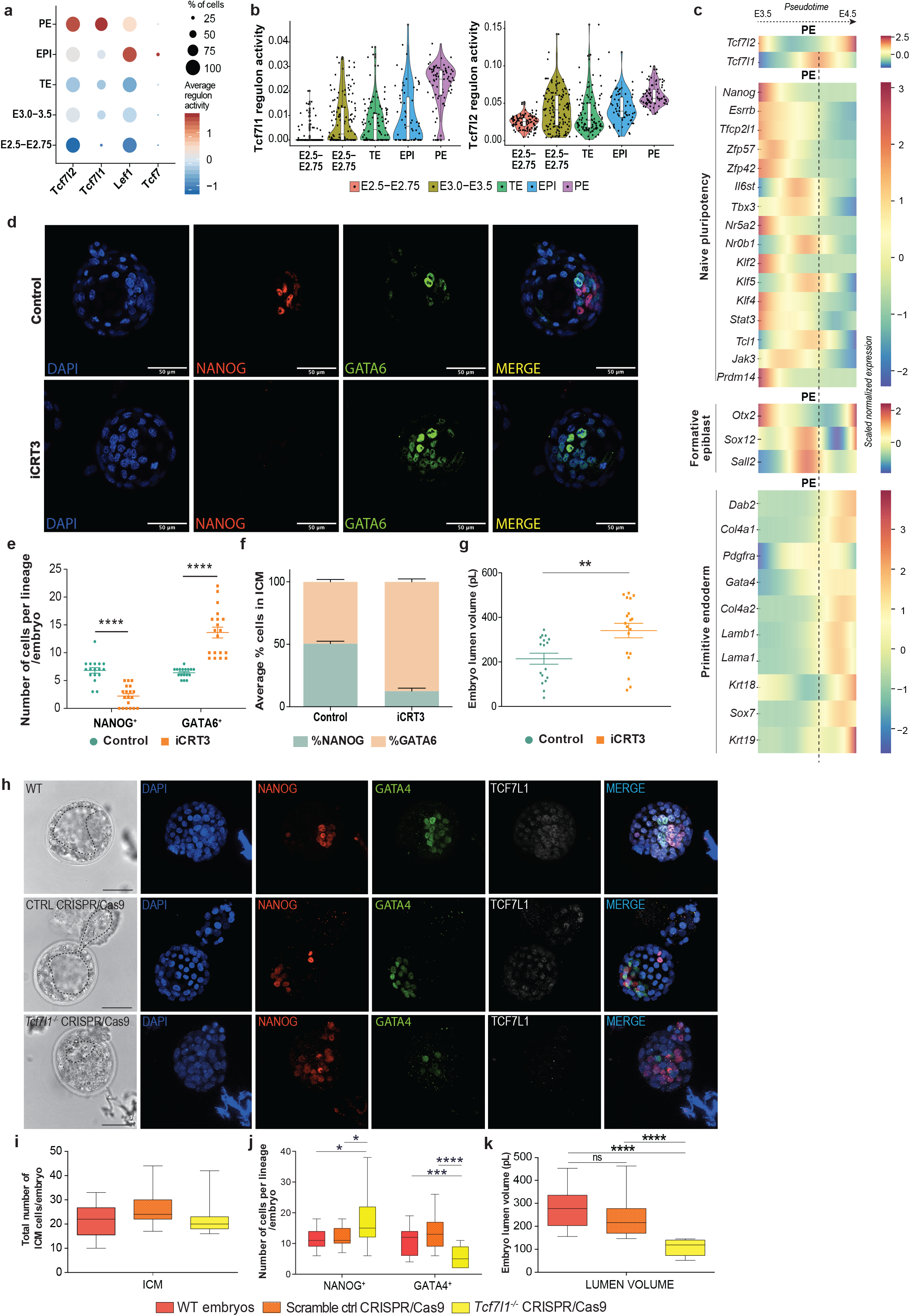
TCF7L1 as main regulator of *in vivo* PE cell fate specification. (a) Regulon activity analysis of TCF/LEF factors during embryo preimplantation development ^87^. Circle size indicates % of cells in which TCF/LEF regulons are active. (b) Violin plot depicting Tcf7l1 and Tcf7l2 regulon activity at different developmental stage of embryo development ^87^. (c) *Tcf7l1* and *Tcf7l2* gene expression along pseudo-time of PE specification trajectory during the transition between E3.5 to E4.5 developmental stages, compared to gene expression levels of different lineage markers ^51^. (d) Representative IF image of preimplantation embryos upon iCRT3 treatment (10μM). Nuclei were counterstained with DAPI. Scale bar=50 μM. (e) Number of NANOG+ and GATA6+ cells (counts) per embryo. Mean ± SEM; Control n=17; iCRT3 n=19; 2 independent experiments. *t* test ****p<0.0001. (f) Percentage of NANOG^+^ and GATA6^+^ cells normalized on total number of ICM per embryo. Mean ± SEM; Ctrl n=17, iCRT3 n=19; 2 independent experiments. (g) Embryo lumen volume reported in pL. Mean ± SEM; Ctrl n=17, iCRT3 n=19 embryos; 2 independent experiments; unpaired *t* test **p< 0.01. (h) Representative IF image of WT, negative control CRISPR/Cas9 and *Tcf7l1^-/-^* CRISPR/Cas9 embryos. Nuclei were counterstained with DAPI. Scale bar=50μM. Dashed lines delimitate lumen cavity. (i) Total number of ICM cells in WT, negative control CRISPR/Cas9 and *Tcf7l1^-/-^* CRISPR/Cas9 embryos. Whiskers represent Min to Max; WT n=15, negative control CRISPR/Cas9 n=19, *Tcf7l1^-/-^* CRISPR/Cas9 n=15. 3 independent experiments. No statistically significant differences using One-way ANOVA test. (j) Number of NANOG+ and GATA4+ cells (counts) per embryo in WT, negative control CRISPR/Cas9 and Tcf7l1-/- CRISPR/Cas9 embryos. Whiskers represent Min to Max; WT n=15, negative control CRISPR/Cas9 n=19, Tcf7l1-/- CRISPR/Cas9 n=15. 3 independent experiments. One-Way ANOVA *p<0.05, ***p<0.001, ****p<0.0001. (k) Embryo lumen volume reported in pL. in WT, negative control CRISPR/Cas9 and *Tcf7l1^-/-^* CRISPR/Cas9 embryos. Whiskers represent Min to Max; WT n=15, negative control CRISPR/Cas9 n=19, *Tcf7l1^-/-^* CRISPR/Cas9 n=15. 3 independent experiments. One-Way ANOVA ****p<0.0001.

Next, we used CRISPR/Cas9-mediated genome editing to further delineate the role of TCF7L1 in regulating cell lineage specification during preimplantation development (Fig. 6h). *Tcf7l1-* targeting ribonucleoproteins (RNPs) were microinjected at the zygote stage (E0.5), resulting in complete editing of the *Tcf7l1* locus, caused mainly by out-of-frame indel mutations (Supplementary Table 10). Only embryos with a 100% efficiency on *Tcf7l1* genome editing were considered for analysis in the *Tcf7l1*-CRISPR/Cas9 group (Supplementary Fig. 6b, c). Control groups consisted of negative control RNP-injected zygotes and WT control zygotes. No difference in the total number of ICM cells was seen between *Tcf7l1^-/-^* embryos and controls (Fig. 6i). However, we observed a profound shift in PE towards EPI cell fate specification in *Tcf7l1^-/-^* embryo ICMs, that now contained a significantly greater number of NANOG^+^ (EPI) cells and a reduced number of GATA4^+^ (PE) (Fig. 6j). Also, the *Tcf7l1^-/-^* blastocyst lumen volume was substantially smaller compared to control embryos (Fig. 6k).

Altogether this study reveals a previously undescribed TCF7L1-dependency in PE formation during preimplantation development. Absence of TCF7L1 induces accumulation of EPI (NANOG^+^) cells with a significative reduction of PE (GATA4^+^) cells.

## Discussion

In this study we investigated the role of canonical Wnt inhibition in preimplantation development and mESC fate *in vitro*. We showed that extrinsic inhibition of Wnt by DKK1 treatment or by transcriptional inhibition (iCRT3) drives PE formation during preimplantation development and mESC cell fate commitment. Furthermore, forced expression of the Wnt-transcriptional repressor, *Tcf7l1*, is sufficient to induce differentiation of mESCs into PE-like cells, but not into EPI-derived lineages. Conversely, ablation of *Tcf7l1* expression diminishes PE formation during preimplantation development and in mESCs lineage commitment without compromising EPI-priming.

The co-emergence of EPI and PE lineages within the ICM of the preimplantation blastocyst requires initial co-expression of the lineage-specific markers NANOG and GATA6 ^4^. The ability of NANOG and GATA6 to antagonize each other ^14,88,89^ suggests that either activation or repression of any of the respective lineage-transcriptional programs may provide the basis for the binary EPI or PE cell fate specification. However, the initial developmental cues that drive the expression of different transcriptional regulators, linked with the acquisition of EPI or PE fate *in vivo*, as well as the PE lineage conversion in mESCs, remain unclear.

Although the mechanisms that govern preimplantation lineage segregation require future investigation, this work already highlights TCF7L1 as part of the initial heterogeneity in the ICM and as a potential candidate for driving PE specification. Notably, TCF7L1 is an intrinsically expressed TF in naive pluripotent mESCs ^4,62^ and in the preimplantation embryo. Pseudotime trajectory generated from published scRNA-seq data on mouse embryos revealed progressive upregulation of PE-specific genes following the “pulse” of *Tcf7l1* expression. This agrees with our RNA-seq data on mESCs where PE gene transcription was induced upon *Tcf7l1* overexpression, underlying the similarity with the *in vivo* expression patterns and highlighting the role of TCF7L1 as a rheostat and a pioneer component of binary EPI vs. PE lineage decisions. The integrated time-series RNA-seq and ChIP-seq data presented here revealed that TCF7L1 binds and represses genes related to formative transition, a prerequisite state prior to primed EPI specification, while we did not identify binding of TCF7L1 on PE promoters. In fact, upon transcriptional Wnt inhibition by *Tcf7l1* overexpression, TCF7L1 dominates and represses embryonic lineage-determining TFs, allowing PE program to take over and to initiate PE specification.

mESCs resemble the embryonic EPI lineage and are strongly biased towards embryonic fates. This is also reflected by the accumulation of repressive chromatin marks on a subset of promoters, including the PE *Gata6* promoter ^6,90^. Nonetheless, previous studies have shown that mESCs have also the potential to transit spontaneously towards an extraembryonic PE-like state ^12–16^. Naive pluripotency is supported by the combined modulation of two cascades, the activation of Wnt and the inhibition of Erk ^21^. Interestingly, MAPK/Erk activation has been reported to regulate binary decisions and force cells to adopt a PE fate in the ICM ^91^. In detail, FGF/MAPK is required for the maintenance of PE cells in the ICM as a result of the inverse expression of Fgf4/Fgfr2 in EPI-fated and PE-fated cells respectively ^23,92^. A later model integrated the FGF signals with the transcriptional changes occurring in EPI/PE cells and suggested that the FGF/MAPK signaling input represses the EPI-specific gene expression program causing the mutually repressive interactions between EPI- and PE-like program ^6^. However, another study highlighted that FGF/MAPK signaling has an additional positive input onto GATA6 expression, and identified positive autoregulatory feedback loops elicited from both NANOG and GATA6 ^93^. Equivalent to embryos, *in vitro* experiments have demonstrated that Erk activation induces PE priming in naive mESCs ^94^. Our findings support that, similarly to MAPK/Erk activation ^94^, Wnt inhibition promotes extraembryonic PE differentiation but not differentiation towards embryonic epiblast-derived lineages. Altogether, these results insinuate that maintenance of ground-state pluripotency conditions by both Erk and Wnt signaling modulation relies on the suppression of PE differentiation rather than the active induction of a naïve pluripotent state.

Although it is well known that the canonical Wnt pathway is indispensable for peri- and post-implantation development ^49^, its role in preimplantation stages remains unclear. Expression of a stabilized isoform of β-catenin ^95^, deletion of *β-catenin* ^96^, *Apc* loss-of-function ^97^ or deletion of *Tcf7l1* ^98^, all affect development progression at post-implantation stages in mouse embryos; however, effects on EPI vs. PE specification were not studied in these cases. Interestingly, *Porcn^-/-^* embryos, which are defective for Wnt secretion, did not show any defect on cell number or cell fate decisions in preimplantation development, indicating that autonomous embryo Wnt ligand secretion is not a prerequisite at these stages. However, Wnt ligands required for canonical and non-canonical cascade activation are expressed in murine and human endometrium in unique patterns, indicative that embryo development does not solely rely on autologous Wnt signals ^53,99,100^. In support of this, our results show that extrinsic inhibition of the Wnt pathway leads in a significant increase of PE cells and reduced EPI cell number, suggesting that the Wnt pathway is important for EPI and PE lineage segregation. We speculate that the post-implantation defects caused by Wnt component deletion might originate from the defective EPI vs. PE segregation during preimplantation development.

Based on *in vitro* experiments, it was previously considered that ablation of *β-catenin* had little or no effect on the self-renewal and transcriptomic signature of mESC ^57,101^. However, a recent study showed that complete deletion of *β-catenin* slightly increases the expression of PE markers, while maintaining the self-renewal state ^102^. We found that *Tcf7l1* overexpression promotes a stronger phenotype than *β-catenin* deletion as it is sufficient to induce EPI to PE lineage conversion, suggesting that the transcriptional inhibitory force of TCF7L1 prevails over the transcriptional effect exerted by β-catenin ^43,63^.

Although our results demonstrate a direct repression of formative genes by TCF7L1 in pluripotent culture conditions, it has been shown that upon naive medium withdrawal, TCF7L1 facilitates formative transition by restraining naive pluripotency network ^8^. However, in agreement with previous studies ^8,61,103^, our results show that deletion of *Tcf7l1* alone delayed but did not abrogate pluripotency exit, EpiSCs commitment or neuroectodermal differentiation. Strikingly, extraembryonic PE differentiation was severely compromised in *Tcf7l1^-/-^* cells. In consistence, only the triple KO of *Tcf7l1* in combination with the transcriptional regulators ETV5 and RBPJ was effective in arresting the cells in the pluripotent state, rendering them refractory to differentiation ^8^. Interestingly, it has been shown that TCF7L1 can repress definitive endoderm genes and its deletion facilitates endoderm specification at initial stages, when cells are cultured in chemically defined endodermal differentiation medium ^104^. This is not in conflict with our results since TCF7L1, as all TCF factors, elicits diverse transcriptional programs in a context- and cell state-dependent manner ^105^. For instance, TCF7L2 has been described as the main driver of metabolic zonation of the liver by activating zonal transcription, in contrast to its reported role as transcriptional repressor in mESCs ^106^.

Here, we provide further evidence demonstrating that distinct TCF factors may regulate independent cellular functions. We demonstrate that deletion or overexpression of *Tcf7l1* has drastic effects on the capacity of mESCs to undergo PE cell lineage conversion, differently from *Tcf7*, which shows not significant effects. We and others have previously shown that the Wnt transcriptional repressor TCF7L1 and the activator TCF7 display unique DNA binding sites ^37^ leading to the regulation of distinct gene sets in mESCs ^37,107^. Explicitly, TCF7 regulates Wnt-dependent cell cycle events by directly binding to cyclin-dependent kinase inhibitors such as *p16Ink4a* and *p19Arf* ^37^. In contrast, we are not able to detect binding of TCF7L1 on cell cycle genes while it is present on genes regulating naïve and formative pluripotency.

In conclusion, our study unravels new aspects of the mechanism governing EPI vs. PE binary cell-fate decisions as part of the interconnected cascades and gene regulatory networks. Further understanding of TCF7L1 function at single-cell and chromatin accessibility level, will contribute to elucidating the complex circuit of differentiation decisions in the preimplantation embryo and beyond.

## Methods

### Cell culture

Undifferentiated wild-type (WT) murine ESCs, *Tcf7l1^-/-^* mESCs obtained from B. Merril ^108^ and *Tcf7^-/-^* mESCs previously generated in the laboratory ^37^ were cultured at 37°C and 5% CO_2_ on gelatin-coated plates in DMEM (Gibco), 15% fetal bovine serum, 2mM L-glutamine (Gibco), 1X minimal essential medium non-essential amino acids, 1x sodium pyruvate, 1x penicillinstreptomycin, 100 μM β-Mercaptoethanol and 1000 U/ml recombinant murine LIF (Peprotech).

For the evaluation of naive EPI and PE-like populations in Fig. 1a; Fig. 3c,d and Supplementary Fig.3b, cells were maintained for at least 3 passages in knockout DMEM (Gibco), 20% knockout serum replacement (KOSR), 2 mM L-glutamine, 1x minimal essential medium non-essential amino acids (Gibco), 1x penicillin/streptomycin (Gibco), 100 μM β-Mercaptoethanol and 1000 U/mL recombinant murine LIF.

For the pharmacological modulation of Wnt pathway in Fig.2a and Fig. 3a, undifferentiated WT ESCs were seeded at a density of 250.000 cells per well in 6-well plates and treated with 200ng/mL Wnt antagonist Dickkopf 1 (DKK1) (Peprotech) or 10μM iCRT3 [inhibitor of β-catenin–responsive transcription] (Sigma) for 3 passages in KOSR/LIF conditions.

### Retinoic acid differentiation

Retinoic acid (RA) differentiation experiments were performed as previously described in Semrau et al., 2017. Explicitly, prior to differentiation cells were grown for at least 2 passages in 2i medium plus LIF (2i/L): DMEM/F12 (Gibco) supplemented with 0.5x N2 supplement, 0.5x B27 supplement, 0.5mM L-glutamine, 20 μg/ml human insulin (Sigma-Aldrich), 1 × 100U/ml penicillin/streptomycin, 0.5x MEM Non-Essential Amino Acids, 0.1 mM 2-Mercaptoethanol, 1 μM MEK inhibitor (PD0325901, Sigma-Aldrich), 3 μM GSK3 inhibitor (CHIR99021, Sigma-Aldrich), 1000 U/ml mouse LIF (Peprotech). Cells were seeded at a density of 2.5 x 10^5^ per 10cm dish and grown over night (12h). The next day cells were washed twice with PBS and medium was replaced with basal N2B27 medium (2i/L medium without inhibitors, LIF and the additional insulin) supplemented with 0.25μM all-trans retinoic acid (Sigma-Aldrich). Medium was being refreshed every 48H.

### *Tcf7l1* transgene induction

*Tcf7l1*-inducible ESC line was purchased from the NIA Mouse ES Cell Bank ^69^. ES cells were cultured on feeder cells in ESC medium containing 0.2μg/ml doxycycline (Dox) (Sigma). Before transgene induction cells were cultured on gelatin coated dish in medium containing 0.2μg/ml Dox for 3 days. One day before transgene induction, 1×10^6^ ES cells were plated onto gelatin coated 100cm^2^ dishes for 24 hours. Dox was removed by washing the cells three times with PBS at interval of 3 hours. In the absence of Dox, the recombinant locus expresses TCF7L1 and Venus YFP protein connected via a synthetic internal ribosomal entry site (IRES). Following the 24h of induction (Day 1), cells were dissociated using 0,5% trypsin-EDTA and resuspended in FACS buffer [PBS supplemented with 5% FBS]. Venus^high^ expressing cells were sorted by fluorescence-activated cell sorting (FACS) using a BD FACSAria™ III sorter and replated in ESC medium conditions. Cells were harvested after 2, 4 and 8 Days after Dox removal.

### EpiLCs differentiation

WT and *Tcf7l1^-/-^* mESCs were maintained for at least 5 passages in N2B27 plus 2i/LIF. Cells were plated at a density of 1×10^4^/cm^2^ in fibronectin (Millipore) coated (16.7 mg/ml) 6-well plates in N2B27 basal medium supplemented with 20ng/ml activin A (Peprotech), 12.5ng/ml Fgf2 (Peprotech) and 2μM XAV939 (Sigma) for 2 and 3 days. For the TCF7L1 overexpressing cells, transgene induction was performed as explained before and Venus^high^ expressing cells were FACS-sorted and replated in the same Epi-inducing medium for 2 and 3 days.

### Mouse Embryo Recovery, Culture and pharmacological modulation of Wnt signaling

To obtain preimplantation embryos, CD-1 female mice were superovulated (SO) by intraperitoneal injection of 150 μL pregnant mare’s serum (PMS) gonadotropin, followed by injection of 150 μL human chorionic gonadotropin (hCG) 48 hours later. SO females were then mated with male mice. For β-CATENIN quantification, E3.5 and E4.5 embryos were flushed from dissected uteri with EmbryoMax® M2 Medium (Sigma). Embryos were washed with PBS and fixed in 4% paraformaldehyde (PFA). For *ex vivo* embryo culture and treatment, E2.5 embryos were flushed from dissected oviducts using EmbryoMax® M2 Medium. E2.5 embryos were treated with 100ng/mL DKK1 and 10μM iCRT3 diluted in EmbryoMax® KSOM Medium and cultured in cell culture dish (Nunc™ Cell-Culture Treated Multidishes, Thermo Fisher Scientific) at 37°C in incubator supplied with 5% CO_2_. Blastocysts (E2.5+48H) were stopped and fixed in 4% PFA for further molecular analysis (see Immunofluorescence staining section).

### Antibody staining and Flow cytometry

For the evaluation of naive EPI and PE-like populations in Fig. 1a; Fig. 3c,d and Supplementary Fig. 3b, cells growing as described above were washed once with PBS and then incubated with Cell dissociation solution (Sigma) for 20 min at 37°C. Cells were washed twice with PBS and were counted. Conjugated antibodies were added in PBS (0,2μg for each 1×10^6^ cells) and samples were incubated for 30 min at 4°C. We used the following antibodies: PE anti-mouse CD140a (PDGFRA) (Thermo Fisher Scientific, 12-1401-81), APC anti-mouse CD31 (PECAM-1) (Thermo Fisher Scientific, 17-0311-82), PE Rat IgG2α (BD Biosciences, 553930), APC Rat IgG2α (Thermo Fisher Scientific, 553930). Following antibody incubation, the cells were washed once in FACS buffer, resuspended in fresh FACS buffer, and passed through filter. For the differentiation experiments shown in Fig. 3e-l and Supplementary Fig.3c-e, cells growing as described above were first dissociated with Cell dissociation solution (Sigma) for 20 min at 37°C. Then, cells were incubated in a volume of 250μl of basal (N2B27) medium with conjugated antibodies (0,2μg for each 1×10^6^ cells) for 30 min at 37°C, in the dark. Afterwards, cells were washed once with PBS, resuspended in basal medium and filtered. Flow cytometry was performed using a BD Canto HTS. Unstained and isotype control samples were used for gating on forward and side scatter. Data analysis was carried out using FlowJo software.

### Gene expression analysis

Total RNA was extracted from cells using GenElute™ Mammalian Total RNA Miniprep Kit. cDNA was reverse transcribed from 500ng of RNA using the iScript cDNA synthesis kit according to manufacturer’s guidelines. Real-time quantitative PCR (RT-qPCR) was performed in three technical replicates per sample using SYBR Green master mix on a ViiA 7 Real-Time PCR system (AB applied biosystems) utilizing specific primers at a concentration of 1μM. Primer sequences used in this study are specified in Supplementary Table 11. Data analysis was performed with QuantStudio™ Real-Time PCR Software. Ct values detected for each sample were normalized to the housekeeping gene *Gapdh*.

### Western blot analysis

ES cells were washed with PBS, trypsinized, and collected by centrifugation. Whole cell lysates were prepared using RIPA cell lysis buffer (Sigma) containing 1:100 phosphatase inhibitor cocktail 2, phosphatase inhibitor cocktail 3 and protease inhibitor (Sigma). Samples were rotated for 30 min at 4°C and spun at max speed for 10 min at 4°C. The supernatant from samples was collected and protein concentration was determined by Bradford assay. Equal amounts of protein per sample were combined with Laemmli buffer, denatured for 5 min at 95°C and subjected to SDS/PAGE separation, followed by immunoblotting. The following primary antibodies were used: rabbit anti-active β-Catenin (Cell Signaling Technology, 8814), mouse anti-total β-Catenin (BD Biosciences, 610154), rabbit monoclonal anti-Tcf1 (Cell Signaling Technology, 2203), rabbit monoclonal anti-Lef1 (Cell Signaling Technology, 2230). Mouse anti-β-ACTIN (Santa Cruz Biotechnology; sc-47778) was used as a load control. Protein quantification was performed with ImageJ software from n=3 independent biological replicates (Fig. 1e).

### Immunofluorescence staining

WT and *Tcf7l1*^-/-^ cells were cultured on gelatin-coated glass coverslips for 4 days as described in the Retinoic acid differentiation section.. *Tcf7l1*-OE Day0, *Tcf7l1*-OE Day6, XEN cells and embryos were cultured as described above. Cells and embryos were washed 2x in PBS, fixed in 4% PFA for 20 min at room temperature (RT) and permeabilized with 0,2% Triton X-100 in PBS/donkey serum (DS) 3% for 10 min. Samples were then blocked with PBS/DS 5% + 0,2% Tween20 + 0,2% BSA for 30 minutes at RT and incubated with primary antibodies overnight (o/n) at 4°C. Primary antibodies were diluted in PBS/DS 3% + 0,02% Tween20 at the specific working concentration. Rat anti-Nanog (eBioscience™, 14-5761-80) (1:200), goat anti-Gata6 (R&D systems, AF1700) (1:200), goat anti-Nestin (Santa Cruz Biotechnology, sc-21248) (1:200), mouse anti-Gata4 (BD Biosciences, 560327) (1:250), rabbit anti-active β-Catenin (Cell Signaling Technology, 8814) (1:200), mouse anti-total β-Catenin (BD Biosciences, 610154) (1:200), goat anti-Tcf3 (Santa Cruz Biotechnology, sc-8635) (Tcf7l1) (1:200). Next, samples were repeatedly washed and incubated with secondary-AlexaFluor antibodies (1:500 in PBS/DS 3% + 0,02% Tween) for 1 hour at RT. DAPI was used to stain nuclei. Lastly, stained embryos were mounted in 10μL PBS/DS 3% drops covered with mineral oil on 35mm glass-bottomed dishes. Embryos were imaged under a Leica SP8x confocal microscope. WT and *Tcf7l1^-/-^* cells were mounted on microscope slides imaged under a Zeiss AxioImager Z1 Microscope using the AxioVision SE64 software. *Tcf7l1*-OE Day0, *Tcf7l1*-OE Day6, XEN cells were imaged under an Operetta CLS™high-content analysis system. Image quantification and analysis was performed using ImageJ software and Harmony High-Content Imaging and Analysis software.

### Chimera generation

To tag the *Tcf7l1*-inducible ESC line (see above) with a constitutively expressed fluorescent protein before embryo injection, a PiggyBac transposase-encoding plasmid (1μg) and pCAG-dsRED-hygro^R^ (200ng) were co-transfected to randomly insert dsRED-hygro^R^ in the genome. Transfection was performed using a Lipofectamine 3000 kit (Invitrogen). Cells were FACS-sorted and kept under the respective drug selection on hygro^R^ feeder cells in Dox-containing medium (0.2μg/ml). To avoid embryo toxicity due to hygromycin, the drug was removed from the culture several passages before injection. Transgene induction was performed as described above (*Tcf7l1* transgene induction section). Following 4 days of induction, cells were dissociated using 0,5% trypsin-EDTA and resuspended in FACS buffer. Venus^high^/dsRed^+^ were sorted using a BD FACSAria™ III sorter and used for blastocyst injections. CD1 embryos were recovered from SO CD1 females at E3.5 and injected with 10-12 Venus^high^/dsRed^+^ cells in EmbryoMax® M2 Medium at the CBD Mouse Expertise Unit of KU Leuven. After injection, blastocysts were cultured for 1-2 hours in EmbryoMax® KSOM Medium at 37°C and 5% CO_2_ and transferred into the uterus of pseudopregnant CD1 females. At E6.5, females were sacrificed using cervical dislocation and embryos were isolated in PBS using a stereo dissection microscope. Embryos were imaged using a Nikon Ti2 eclipse microscope.

### Whole immunofluorescence staining

Dissected E6.5 stage embryos were fixed for 1 hour in 4% PFA. Staining was performed as described in Kalkan et al., 2019. Primary goat anti-Gata6 (R&D systems, AF1700) (1:200) and rabbit anti-dsRed (Takara, 632496) (1:200) antibodies were used to mark visceral endoderm and trace dsRed^+^ cells respectively. Secondary-AlexaFluor antibodies were used to detect primary antibodies. Embryos were mounted in 50% glycerol in PBS on 35mm glass-bottomed dishes and were imaged using a Leica SP8x confocal microscope. Images were processed with Fiji software.

### Sample preparation and RNA sequencing

RNA extraction was performed using GenElute™ Mammalian Total RNA Miniprep Kit using On-Column DNase I Digestion Set. RNA was quantified with Nanodrop 1000 spectrophotometer (Thermo Fisher Scientific) and analyzed using a 2100 Bioanalyzer (Agilent Technologies). Three biological replicates were prepared for each sample. Libraries were produced using KAPA Stranded RNA-Seq Library Preparation Kit Illumina® Platforms. Final cDNA libraries were checked for quality using a 2100 Bioanalyzer and quantified using Qubit dsDNA HS Assay Kit. The libraries were loaded in the flow cell at 4nM concentration and sequenced on the Illumina HiSeq4000 sequencing system producing single-end 50-nucleotide reads at Genomics Core Leuven (Belgium).

### Quality assessment and Mapping

Adapters were removed using Cutadapt and processed reads shorter than 20bp were discarded. Mapping was performed with HISAT2 v2.1.0, against the mouse reference genome (mm10) with default parameters. Alignment scores (percentages of uniquely mapped reads) ranged from 71.2% to 81.5%. A grand total of >340Mi uniquely mapped reads was used for further analyses.

### Gene assignment and Differential Gene Expression Analysis

FeatureCounts v2.0.1 ^109^ was used to count mapped reads for genomic features in the mouse transcriptome (Ensembl Release M25, GRCm38.p6). A total of 18584 genes with average count greater than 5 were kept and passed to DESeq2 v1.28.1 ^110^ for the detection of statistically significant differentially expressed genes (Supplementary Table 4; Supplementary Table 5). Differentially expressed genes were called on the basis of DESeq2 thresholds for absolute log2-fold-change (≥2) and a 5% false discovery rate (Fig. 5b-d).

### Functional Enrichment Analysis

Functional enrichment analysis of gene expression was performed with a combination of the Cytoscape ClueGO plug-in and a custom R function based on the igraph library. Differentially expressed gene lists were analyzed with ClueGO. Community detection was applied on the resulting network using the Louvain algorithm, while also taking into account edge betweenness centrality of the functional terms. Network communities were associated to functional terms on the basis of term name over-representations and color-coded accordingly (Fig. 5a; Supplementary Table 2; Fig. 5e; Supplementary Table 3). Gene enrichments at the level of transcriptional regulation were calculated against the TTRUST database using Metascape ^85^ (Supplementary Fig. 5a).

### TCF7L1 binding analysis

Information for TCF7L1 binding was obtained from publicly available ChIP-seq data (E-MTAB-4358) ^37^. TCF7L1 peaks were called using GEM as described in De Jaime-Soguero et al., 2017. Reads were aligned to the latest mouse genome available (mm10, Genome Reference Consortium GRCm38) and genes with TCF7L1 peaks within 10kb around the transcription start site (TSS) were considered to be bound by TCF7L1. Functional enrichment on the intersection of TCF7L1-bound and day1 differentially downregulated genes was performed with gProfiler (Fig. 5h) ^111^. All analyses of gene expression were performed in R. Visualizations of TCF7L1 binding data were made with the use of the IGV Genome Browser ^112^.

### Gene regulatory network analysis using SCENIC

Gene regulatory network analyses in Fig. 6a,b were performed using publicly available resource dataset from Posfai et al., 2021 (GSE145609). Gene regulatory network inference was performed using pySCENIC v0.9.15 ^113^ and regulon activity in samples from Mohammed et al., 2017 dataset (GSE100597) was extracted from the integrated single-cell mouse preimplantation development atlas ^87^. Visualization was performed using DotPlot and and VlnPlot functions in Seurat v3.0.0 ^114^.

### Single-cell RNAseq analysis

Single-cell RNA-seq analyses shown in Fig. 6c, Supplementary 1b,cand Supplementary 6a were performed using publicly available dataset and code as previously described in Nowotschin et al., 2019 (GSE123046). Briefly, raw read counts from two replicate samples of the mouse embryonic development at E3.5 and E4.5 stages were used as an input for Harmony algorithm v0.1.4 ^51^. Raw reads were normalized to the library size and log transformed, highly variable genes were used to construct Harmony augmented affinity matrix which was then used to generate a visualization with a force directed layout (Supplementary Fig. 1b and Supplementary Fig. 1c). The differentiation trajectory was inferred using Palantir algorithm v1.0.0 ^115^ and Harmony augmented affinity matrix. Branches were considered PE or EPI if the corresponding branch probability was higher than 0.9 (Supplementary Fig. 1b; Fig. 6c and Supplementary Fig. 6a).

Gene expression analysis in Fig. 1g and Supplementary Fig. 1e was performed using published datasets ^50,51^. Normalized read counts were used as input for Seurat DoHeatmap function ^114^.

### Ovarian stimulation and zygote collection

B6D2F1 hybrid female mice (6-12 weeks old, Charles River Laboratories, Belgium) underwent ovarian stimulation, by intraperitoneal injection of 5IU pregnant mare’s serum gonadotropin (PMSG, Folligon®, Intervet, Boxmeer, Netherlands), and 5IU human chorionic gonadotrophin (hCG) 48 hours apart, after which natural mating was allowed o/n. Zygotes were removed from the ampulla of the oviduct, followed by 3-minute incubation in 200 IU/ml hyaluronidase, and several washing steps in KSOM medium supplemented with 0.4% bovine serum albumin (Millipore).

### CRISPR/Cas9 zygote microinjection and embryo culture

*Tcf7l1*-targeting CRISPR RNA (crRNA) (protospacer sequence 5’-GGTCTGGAATCATCAGGAAG - 3’, custom, Integrated DNA Technologies, Belgium), negative control crRNA (1072544, Integrated DNA Technologies, Belgium), and transactivating crRNA (tracrRNA) (1072533, Integrated DNA Technologies, Belgium), were dissolved in duplex buffer (1072547, Integrated DNA Technologies, Belgium). A crRNA:tracrRNA complex (guide RNA, gRNA) was formed by mixing the crRNA and tracrRNA in a 1:1 molar ratio, followed by 5 minutes incubation at 95°C. Once the mixture cooled down to room temperature, Cas9 protein (1072545, Integrated DNA Technologies, Belgium) was added to the mixture to form a ribonucleoprotein (RNP), after which the RNP was further diluted in Gamete Buffer, to a final concentration of 25 ng/μl gRNA and 50 ng/μl Cas9, aliquoted and stored at −80°C until further use. Zygotes were piezo-electric microinjected in KSOM medium supplemented with 20% fetal bovine serum (Gibco BRL, Gaithersburg, USA) by aspirating an amount of the RNP complexes, equal to the zygote diameter. Following injection, mouse embryos were cultured until the 8-cell stage in KSOM medium supplemented with 0.4% bovine serum albumin (Millipore), after which the medium was changed to Cook Blasto ® (Cook Ireland Ltd, Ireland). Incubation took place at 37°C, 5% O_2_ and 6% CO_2_ in all cases.

### DNA extraction and sequence analysis

DNA was extracted from single fixed and immune-stained embryos with Arcturus picopure™ DNA extraction kit in a volume of 10 μl extraction solution, according to the manufacturer’s instructions. Extracted DNA was PCR-amplified (F 5’ – GTGCCTTCTCCGTCAGTCTC – 3’, R 5’ – GCAGGCACAAATCCAAGTTT – 3’, and subsequently subjected to next-generation sequencing in an Illumina MiSeq platform ^116^. Raw sequencing data were analyzed by the BATCH-GE tool ^117^.

### Image analysis

Embryo imaging was performed on Leica SP8x confocal microscopy with precise galvo-Z for quick Z-stack imaging. To quantify total and nuclear active (non-phosphorylated) β-Catenin signals, the cells of NANOG- and GATA6-positive cells were manually selected by using ImageJ software from single 2D confocal images. The total active (membrane + cytoplasmatic+ nuclear) and nuclear active (non-phosphorylated) β-Catenin signal levels were measured by ImageJ software. Total active β-Catenin signal in NANOG+ and GATA6+ cells was calculated selecting both membrane and nuclear signals. Nuclear active β-Catenin signal in NANOG+ and GATA+ cells was calculated selecting specifically the DAPI signal area. Embryos were imaged using the same imaging parameters (laser power and smart gain). Images were merged and/or quantified using ImageJ software. Embryo lumen volume measurement of embryo treated with DKK1, iCRT3 and *Tcf7l1^-/-^* was performed by analyzing blastocyst z-stacks obtained using Leica SP8x confocal microscopy with precise galvo-Z for quick Z-stack imaging. The total length of the z-stack was divided by the total number of frames per embryo. Therefore, a constant number representing the height of the z-stack step size was identified using the Leica Application Suite X (LAS X, v3.7.4) software. The lumen area of each frame was calculated using ImageJ. Afterwards, lumen area was multiplied by the height of the z-stack step size. The summary provides the lumen volume of the whole embryo in mm3. Volume was then converted to pico liter (pL) by dividing the lumen volume in mm3 by 1000. Cell fate specification analysis was performed by evaluating lineage-specific transcription factor (TF) signal from each nucleus. Intensity signals from NANOG, GATA6 and GATA4 were normalized on the total number of cells per embryo (DAPI+) and reported as percentage. Detailed differences in cell fate specification were evaluated by dividing the TF signal intensity each marker by the total number of ICM cells for each embryo and reported as percentage.

### Ethical approval

All the experiments performed were approved by the Ethics Committee at KU Leuven University under the ethical approval codes P170/2019 and by Animal Ethics Committee of Ghent University Hospital (approval number ECD 18-29). B6D2F1 mice were obtained from Charles River Laboratories, Brussels, Belgium.

## Statistical Analysis

All statistical analyses in this study were performed using GraphPad Prism 6 software (San Diego, CA, USA) and R Project for Statistical Computing (*R v3.6.1*). Significant differences were determined by One-Way and Two-Way analyses of variance ANOVA (multiple groups) and multiple unpaired t test (two groups). Data are presented as mean ± standard error of mean (SEM) or standard deviation (SD), as Min to Max and as fold change. Statistical differences were indicated as: * p<0,05; ** p< 0,01; *** p< 0,001; **** p< 0,0001.

## Materials Availability

Requests for materials should be addressed to Frederic Lluis and Paraskevi Athanasouli.

## Data Availability

The accession number for the RNA-seq reported in this paper is Gene Expression Omnibus (GEO): GSE171204. Source data are provided with this paper.

## Acknowledgments

We would like to thank Dr. Brad Merrill and for providing WT and *Tcf7l1^-/-^* mESCs cells and Samantha Zaunz for performing the FACS needed for the chimeras generation. The authors would like to thank the VIB Mouse Expertise Unit for the mouse blastocysts injections and the KU Leuven FACS Core for their assistance with the cell sorting. The authors would like to extend their gratitude to the Research Foundation – Flanders for the Ph.D. fellowships awarded to PA (11M7822N), BV (11E7920N), KU Postdoctoral Mandate awarded to ADJS (PDM/18/212) and FWO Research Project Grants G097618N, G091521N (FLL), G073622N (FLL, BH) and C1 KU Leuven internal grants C14/21/115 (FLL).

## Supplementary Figure legends

**Supplementary Fig. 1.**
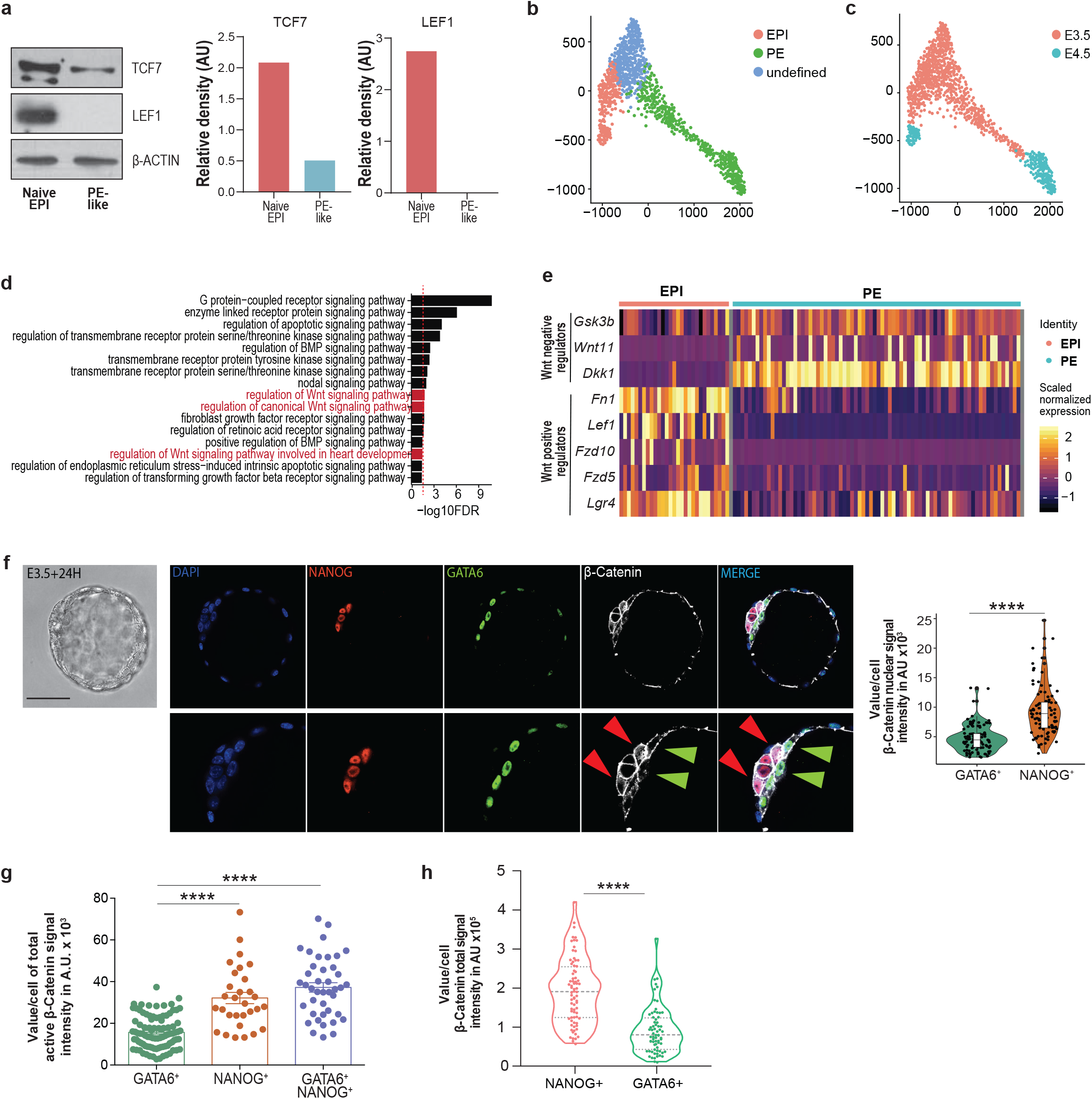
PE cells are characterized by a diminished Wnt/β-Catenin signaling activity compared to the EPI lineage. (a) TCF7 and LEF1 protein levels in naive EPI and PE-like cells. (b) Force-directed layout displaying relationships between undefined ICM, EPI and PE cells. Cells are colored by lineage ^51^. (c) Force-directed layout of E3.5 and E4.5 cells. Cells are colored by embryo developmental timepoint ^51^. (d) GO enrichment analysis of DEGs between EPI and PE lineages in E4.5 preimplantation embryos ^50^. Red dashed line = FDR 0.05. (e) Differential Wnt signaling transcriptome profile between EPI and PE lineages in E4.5 preimplantation embryos ^50^. (f) Representative immunofluorescence (IF) image of active β-catenin protein signal in E3.5+24H *ex vivo* cultured embryos. EPI-NANOG^+^cells (red arrow) and PE-GATA6^+^ cells (green arrow). Scale bar=50 μM. Zoomed region of interest is reported below the images. (g) Total cellular active β-catenin signal intensity in EPI-NANOG^+^, PE-GATA6^+^ and double-positive NANOG^+^/GATA6^+^ cells in E3.5 embryos. Mean ± SEM; NANOG^+^ n=30, GATA6^+^ n=93 and undefined NANOG^+^/GATA6^+^ cells n=41; 2 independent experiments; One-way ANOVA ****p<0.0001. (h) Total active β-catenin signal in EPI-NANOG+ and PE-GATA6+ cells. Integrated intensity in arbitrary units (AU). Mean ± SD; NANOG+ n=76; GATA6+ n=69; 2 independent experiments; t test ****p<0.0001.

**Supplementary Fig. 2.**
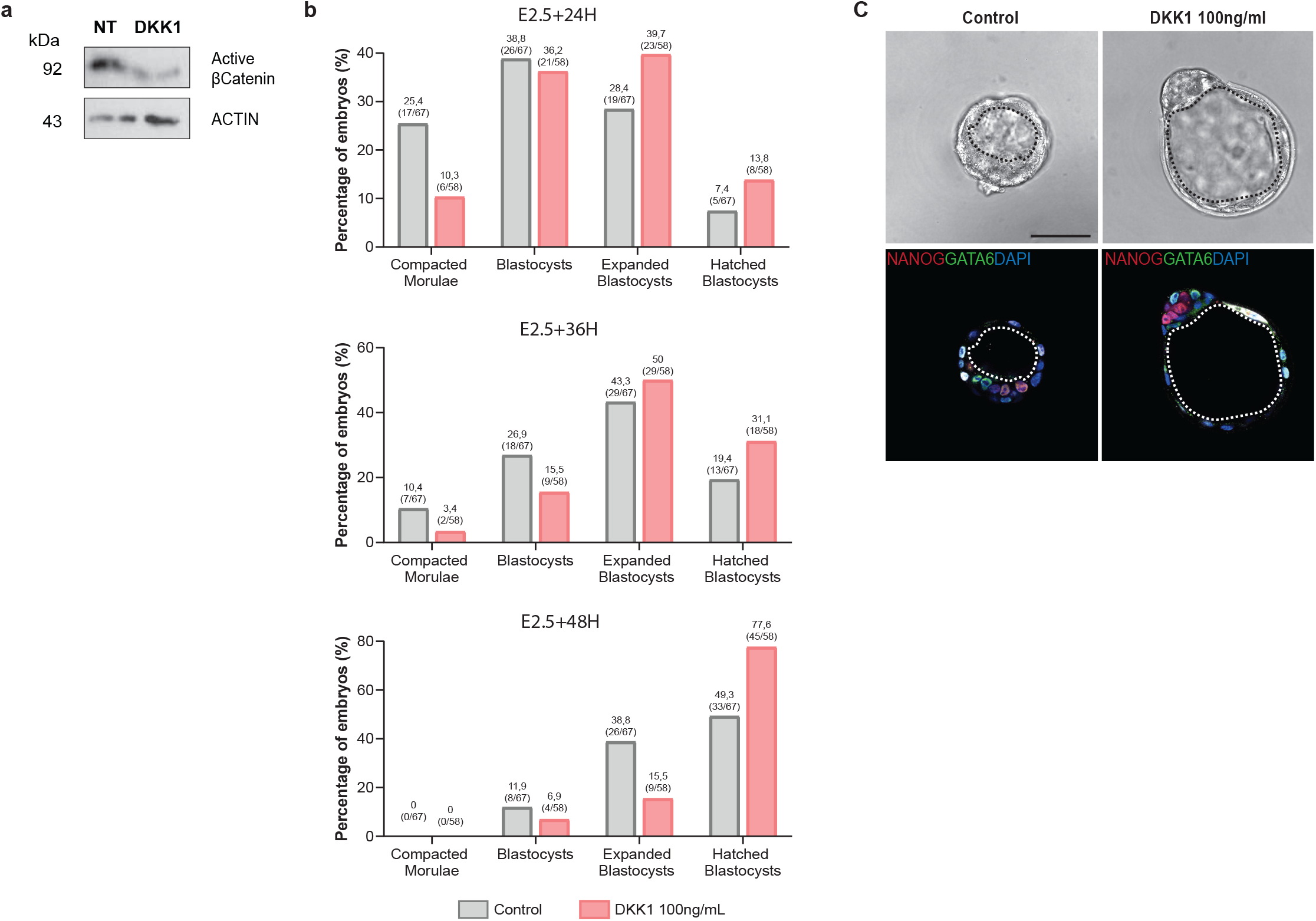
WNT signaling inhibition as major contributor of PE lineage development. **(a)** Representative images of active β-catenin levels in mESCs upon DKK1 treatment for 3 passages. (b) Effect of DKK1 on mouse embryo developmental kinetics. Bars represent percentage of each developmental stage in each treatment group. Combined data of 2 *in vitro* culture sessions. (c) Representative IF image of embryo morphology upon DKK1 treatment. Scale bar = 50μM.

**Supplementary Fig. 3.**
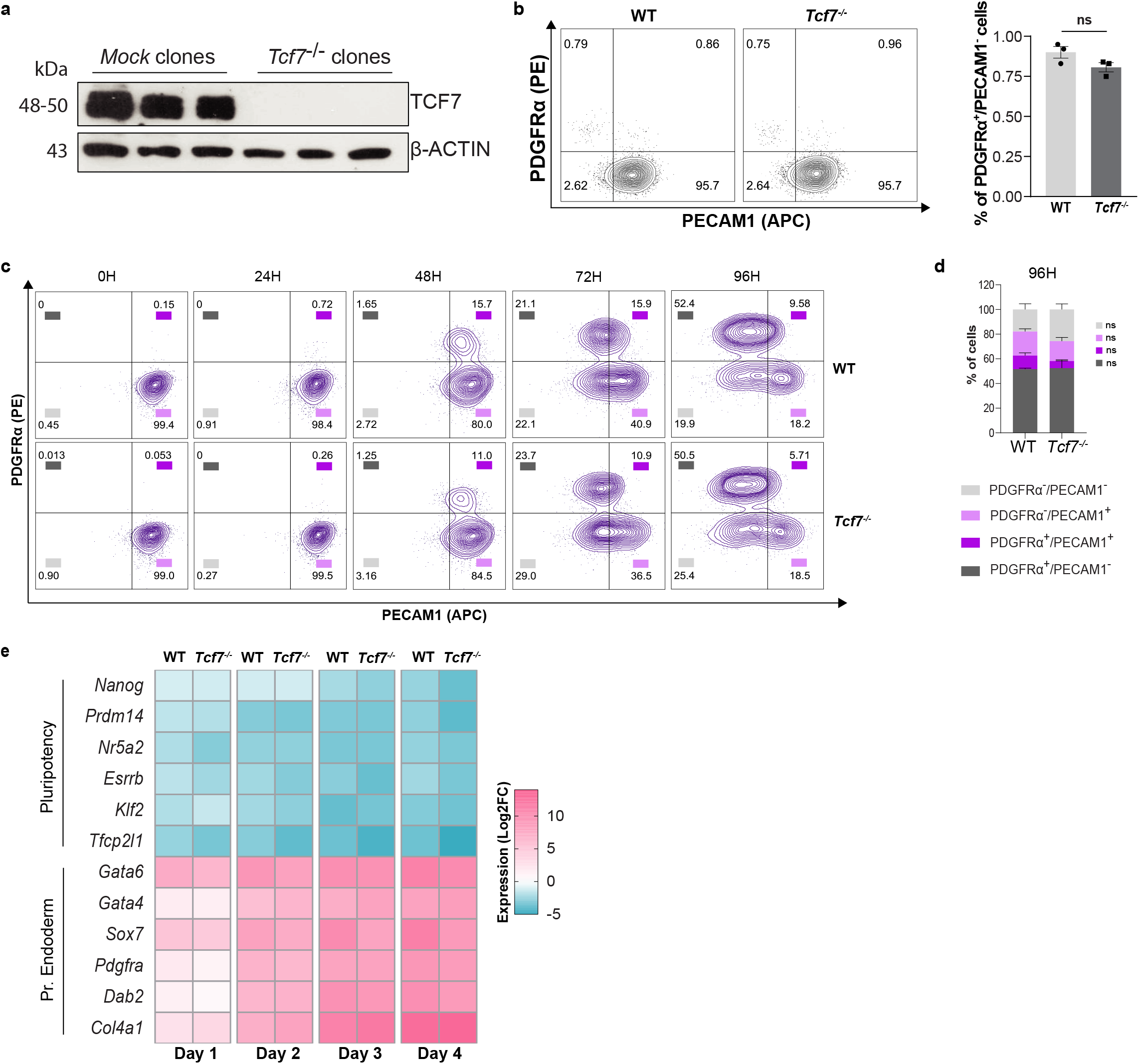
The loss of *Tcf7l1* impedes PE commitment without affecting embryonic lineages specification. (a) TCF7 protein levels in three independent mock and *Tcf7^-/-^* clones. (b) (Left) Representative flow cytometry images of mESC populations co-stained for PECAM1 and PDGFRα in WT and *Tcf7^-/-^* cells. (Right) Flow cytometry analysis of PDGFRα^+^/PECAM1^-^ cells in WT and *Tcf7^-/-^* cells. Mean ± SD; n=3. *t* test. ns=non-significant. (c) Representative flow cytometric images of mESC populations co-stained for PECAM1 and PDGFRα markers in WT and *Tcf7^-/-^* cells upon 0, 24, 48, 72 and 96H of 0.25μM RA treatment. n=3. (d) Flow cytometry analysis of PDGFRα and PECAM1 populations at 96H of RA treatment. Mean ± SD; n=3; two-way ANOVA test. ns= non significant. (e) Gene expression analysis of pluripotency and PE gene markers in WT and *Tcf7*^-/-^ upon RA differentiation. Gene expression values are reported as Log2 of fold change expression.

**Supplementary Fig. 4.**
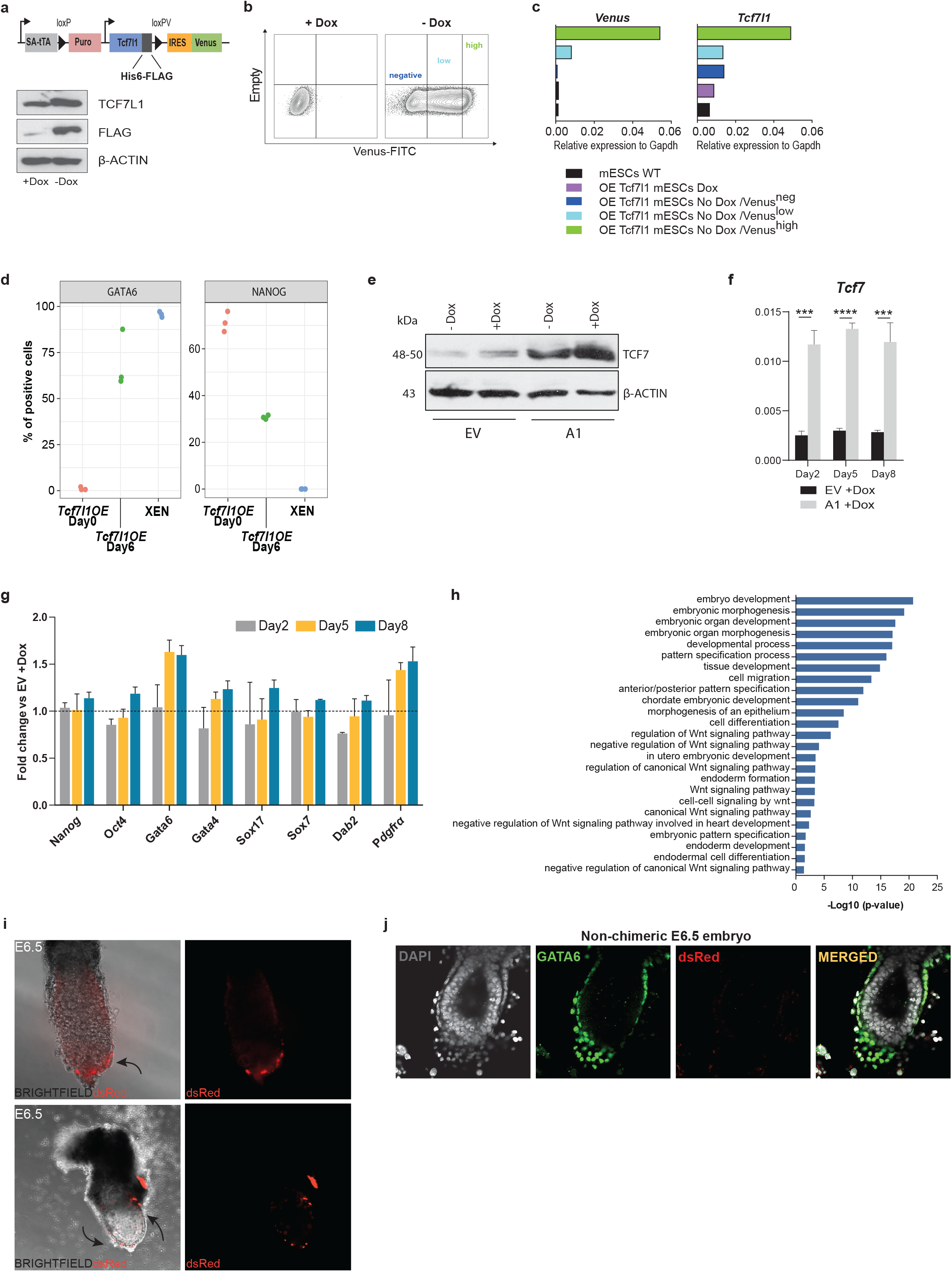
*Tcf7l1* overexpression promotes PE commitment of ESCs. (a) Schematic representation of *Tcf7l1* transgene as designed by Nishiyama et al. 2009. (b) Representative flow cytometry images of Venus expression in uninduced (+Dox) and 24h induced cells (-Dox). Lines separate negative, low and high *Venus*-expressing populations. (c) Gene expression analysis of *Venus* and *Tcf7l1* in sorted cell populations after 24h of transgene induction. Populations were sorted based on *Venus* expression into Venus^neg^, Venus^low^ and Venus^high^. WT mESCs and uninduced (+Dox) cells were used as control. Gene expression was normalized to housekeeping gene *Gapdh*. (d) Percentage of GATA6 and NANOG positive nuclei normalized on the total number of cells (DAPI signal). Each dot represents a field of view. (e) TCF7 levels upon transgene induction with Dox treatment. EV: empty vector, A1: cell clone with stably incorporated transgene cassette. (f) Gene expression analysis of *Tcf7* upon transgene induction for 2, 5 and 8 days. EV treated with Dox was used as control. Mean ± SD; n=3. Multiple *t* tests **p<0.01, ***p<0.001. (g) Gene expression analysis of pluripotency and extraembryonic endoderm genes normalized to *Gapdh* upon *Tcf7* overexpression for 2, 5 and 8 days. Values are reported as fold change relative to EV treated with Dox. (h) GO enrichment analysis of upregulated genes after 8 days of *Tcf7l1 OE*. (i) Representative BF and fluorescence image of E6.5 chimeric embryos injected with *Tcf7l1* induced cells (Venus^high^/dsRed^+^). (j) Non-chimeric E6.5 embryo stained with GATA6 and dsRed. Scale bar=50μM.

**Supplementary Fig. 5.**
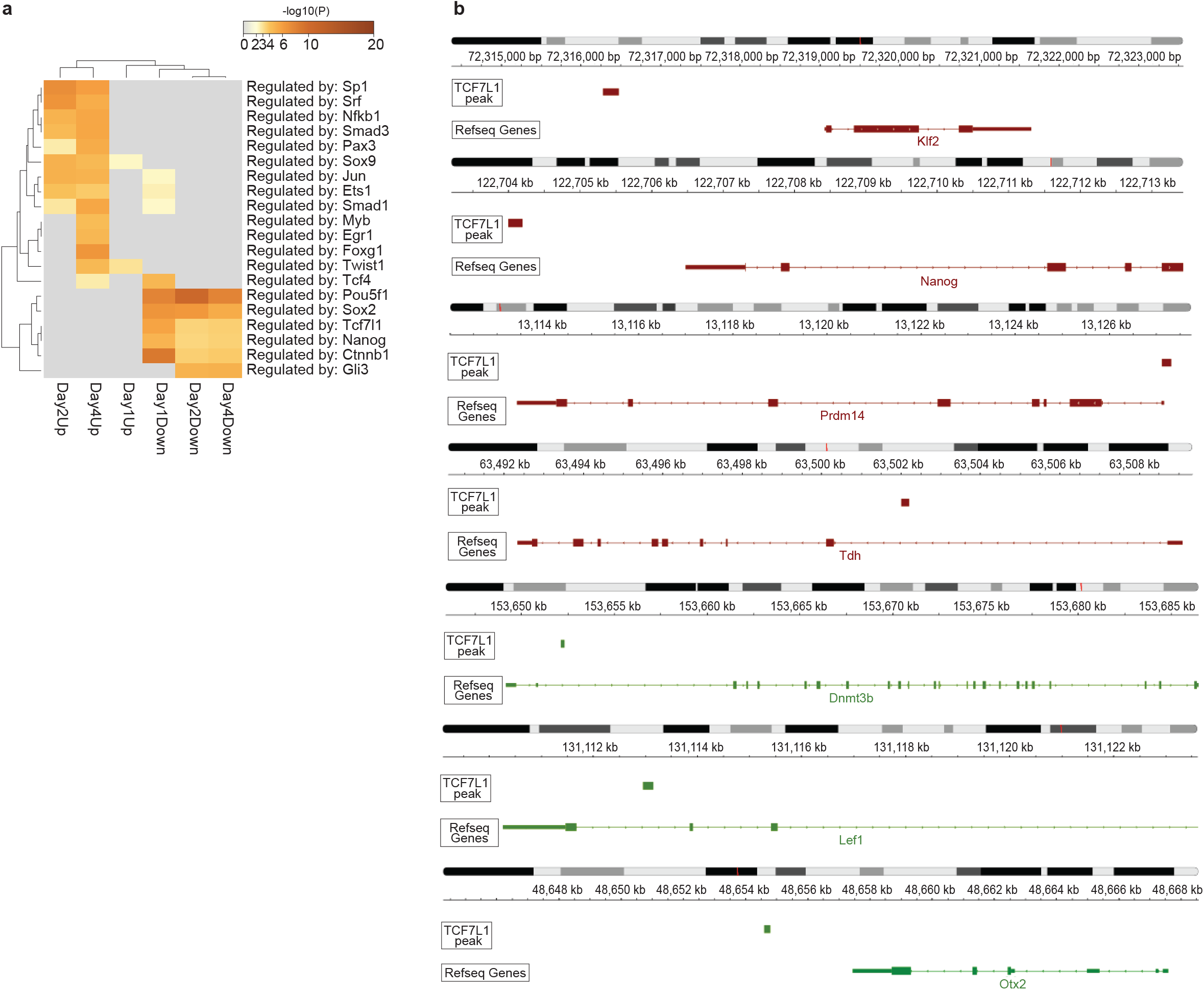
*Tcf7l1* overexpression promotes conversion to PE fate through a downregulation of naive and formative pluripotency. (a) Transcriptional regulator target gene enrichments for genes up- or down-regulated at D1, D2 and D4 of *Tcf7l1* induction. Genes were selected at |logFC|≥2 and adjusted p-value≤0.05. (b) Genomic regions around genes associated with naive, general and formative pluripotency genes with TCF7L1 binding peaks around the TSS region.

**Supplementary Fig. 6.**
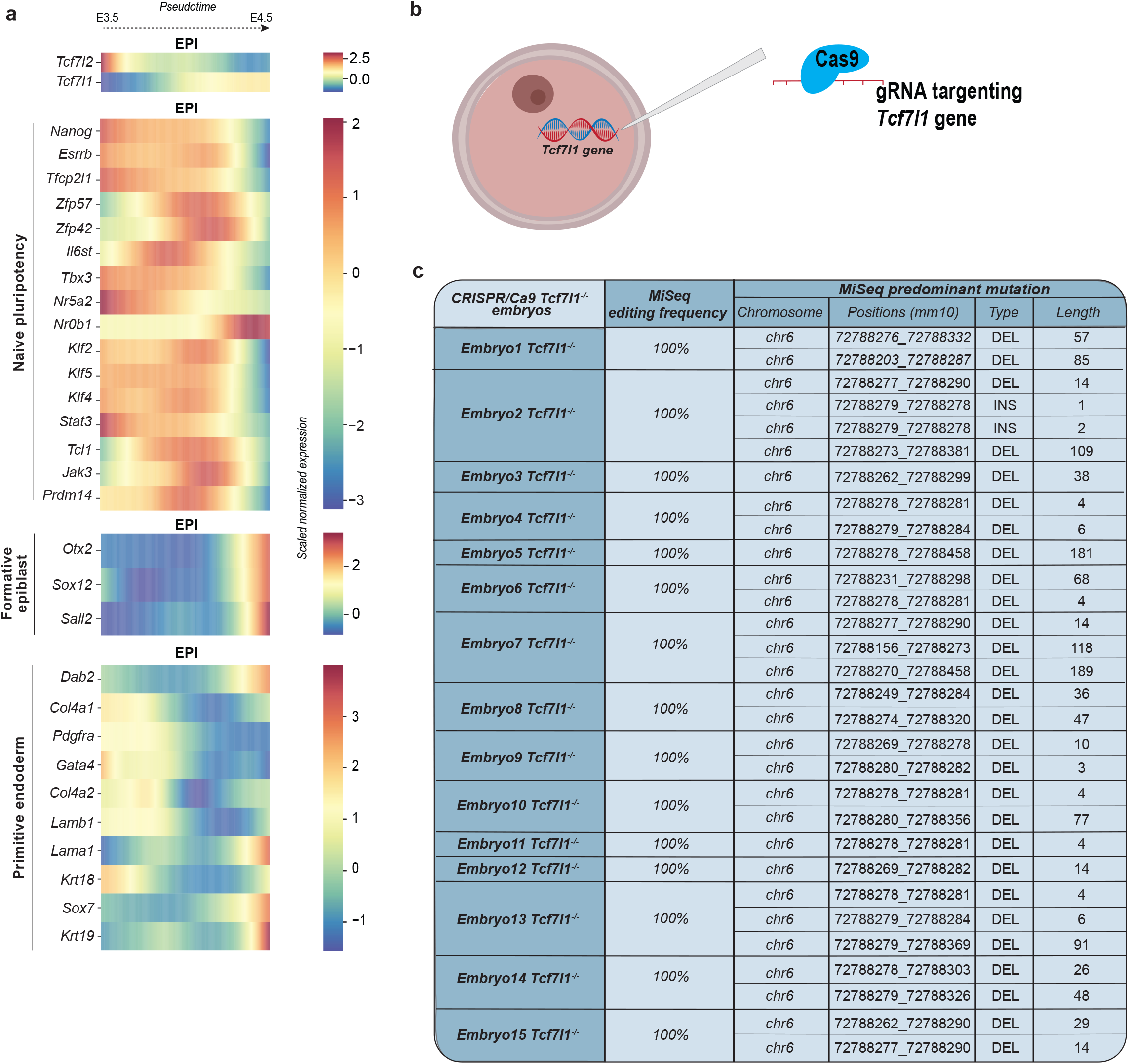
TCF7L1 as main regulator of *in vivo* PE cell fate specification. **(a)** *Tcf7l1* and *Tcf7l2* gene expression profile in EPI lineage from E3.5 to E4.5 compared to gene expression levels of different lineage markers. (b) Representative scheme of CRISPR/Cas9 *in vivo* experiment. (c) MiSeq results of CRISPR/Cas9 *Tcf7l1^-/-^* embryos.

## References

1. Cockburn, K. & Rossant, J. Making the blastocyst: lessons from the mouse. J. Clin. Invest. 120, 995–1003 (2010).

2. Saiz, N. & Plusa, B. Early cell fate decisions in the mouse embryo. Reproduction 145, R65–80 (2013).

3. Hermitte, S. & Chazaud, C. Primitive endoderm differentiation: from specification to epithelium formation. Philos. Trans. R. Soc. B Biol. Sci. 369, 20130537 (2014).

4. Yamanaka, Y., Lanner, F. & Rossant, J. FGF signal-dependent segregation of primitive endoderm and epiblast in the mouse blastocyst. Development 137, 715–724 (2010).

5. Chazaud, C., Yamanaka, Y., Pawson, T. & Rossant, J. Early Lineage Segregation between Epiblast and Primitive Endoderm in Mouse Blastocysts through the Grb2-MAPK Pathway. Dev. Cell 10, 615–624 (2006).

6. Schröter, C., Rué, P., Mackenzie, J. P. & Arias, A. M. FGF/MAPK signaling sets the switching threshold of a bistable circuit controlling cell fate decisions in embryonic stem cells. Dev. 142, 4205–4216 (2015).

7. Plusa, B. & Hadjantonakis, A. K. Embryonic stem cell identity grounded in the embryo. Nat. Cell Biol. 16, 502–4 (2014).

8. Kalkan, T. et al. Complementary Activity of ETV5, RBPJ, and TCF3 Drives Formative Transition from Naive Pluripotency. Cell Stem Cell 24, 785–801.e7 (2019).

9. Kinoshita, M. et al. Capture of Mouse and Human Stem Cells with Features of Formative Pluripotency. Cell Stem Cell 28, 453–471.e8 (2021).

10. Boroviak, T. et al. Lineage-Specific Profiling Delineates the Emergence and Progression of Naive Pluripotency in Mammalian Embryogenesis. Dev. Cell 35, 366–382 (2015).

11. Smith, A. Formative pluripotency: the executive phase in a developmental continuum. Development 144, 365–373 (2017).

12. Morgani, S. M. et al. Totipotent Embryonic Stem Cells Arise in Ground-State Culture Conditions. Cell Rep. 3, 1945–1957 (2013).

13. Lo Nigro, A. et al. PDGFRa+ Cells in Embryonic Stem Cell Cultures Represent the In Vitro Equivalent of the Pre-implantation Primitive Endoderm Precursors. Stem Cell Reports 8, 318–333 (2017).

14. Wamaitha, S. E. et al. Gata6 potently initiates reprograming of pluripotent and differentiated cells to extraembryonic endoderm stem cells. Genes Dev. 29, 1239–1255 (2015).

15. Shimosato, D., Shiki, M. & Niwa, H. Extra-embryonic endoderm cells derived from ES cells induced by GATA Factors acquire the character of XEN cells. BMC Dev. Biol. 7, (2007).

16. McDonald, A. C. H., Biechele, S., Rossant, J. & Stanford, W. L. Sox17-mediated XEN cell conversion identifies dynamic networks controlling cell-fate decisions in embryo-derived stem cells. Cell Rep. 9, 780–93 (2014).

17. Singh, A. M., Hamazaki, T., Hankowski, K. E. & Terada, N. A Heterogeneous Expression Pattern for Nanog in Embryonic Stem Cells. Stem Cells 25, 2534–2542 (2007).

18. Canham, M. A., Sharov, A. A., Ko, M. S. H. & Brickman, J. M. Functional Heterogeneity of Embryonic Stem Cells Revealed through Translational Amplification of an Early Endodermal Transcript. PLoS Biol. 8, (2010).

19. Lanner, F. & Rossant, J. The role of FGF/Erk signaling in pluripotent cells. Development 137, 3351–3360 (2010).

20. Ying, Q. L. et al. The ground state of embryonic stem cell self-renewal. Nature 453, 519–523 (2008).

21. Silva, J. et al. Promotion of reprogramming to ground state pluripotency by signal inhibition. PLoS Biol. 6, 2237–2247 (2008).

22. Morgani, S. M. et al. Totipotent Embryonic Stem Cells Arise in Ground-State Culture Conditions. Cell Rep. 3, (2013).

23. Krawchuk, D., Honma-Yamanaka, N., Anani, S. & Yamanaka, Y. FGF4 is a limiting factor controlling the proportions of primitive endoderm and epiblast in the ICM of the mouse blastocyst. Dev. Biol. 384, 65–71 (2013).

24. Kang, M., Piliszek, A., Artus, J. & Hadjantonakis, A. FGF4 is required for lineage restriction and salt-and-pepper distribution of primitive endoderm factors but not their initial expression in the mouse. Dev. Stem Cells 140, 267–279 (2013).

25. Ohnishi, Y. et al. Cell-to-cell expression variability followed by signal reinforcement progressively segregates early mouse lineages. Nat. Cell Biol. 16, 27–37 (2014).

26. Hamazaki, T., Kehoe, S. M., Nakano, T. & Terada, N. The Grb2/Mek Pathway Represses Nanog in Murine Embryonic Stem Cells. Mol. Cell. Biol. 26, 7539–7549 (2006).

27. Lloyd, S., Fleming, T. P. & Collins, J. E. Expression of Wnt genes during mouse preimplantation development. Gene Expr. patterns 3, 309–12 (2003).

28. Wang, Q. T. et al. A genome-wide study of gene activity reveals developmental signaling pathways in the preimplantation mouse embryo. Dev. Cell 6, 133–144 (2004).

29. Kemp, C., Willems, E., Abdo, S., Lambiv, L. & Leyns, L. Expression of all Wnt genes and their secreted antagonists during mouse blastocyst and postimplantation development. Dev. Dyn. 233, 1064–1075 (2005).

30. Xie, H. et al. Inactivation of nuclear Wnt-β-catenin signaling limits blastocyst competency for implantation. Development 135, 717–727 (2008).

31. Chen, Q. et al. Embryo-uterine cross-talk during implantation: the role of Wnt signaling. Mol. Hum. Reprod. 15, 215–221 (2009).

32. Berge, D. Ten et al. Embryonic stem cells require Wnt proteins to prevent differentiation to epiblast stem cells. Nat. Cell Biol. 13, 1070–1077 (2011).

33. Fan, R. et al. Wnt/Beta-catenin/Esrrb signalling controls the tissue-scale reorganization and maintenance of the pluripotent lineage during murine embryonic diapause. Nat. Commun. 11, 5499 (2020).

34. Cadigan, K. M. Wnt signaling: complexity at the surface. J. Cell Sci. 119, 395–402 (2006).

35. Kohn, A. D. & Moon, R. T. Wnt and calcium signaling: β-Catenin-independent pathways. Cell Calcium 38, 439–446 (2005).

36. Xu, Z. et al. Wnt/β-catenin signaling promotes self-renewal and inhibits the primed state transition in naïve human embryonic stem cells. Proc. Natl. Acad. Sci. U. S. A. 113, E6382–E6390 (2016).

37. De Jaime-Soguero, A. et al. Wnt/Tcf1 pathway restricts embryonic stem cell cycle through activation of the Ink4/Arf locus. PLOS Genet. 13, e1006682 (2017).

38. Cole, M. F., Johnstone, S. E., Newman, J. J., Kagey, M. H. & Young, R. A. Tcf3 is an integral component of the core regulatory circuitry of embryonic stem cells. Genes Dev. 22, 746–755 (2008).

39. Marson, A. et al. Connecting microRNA genes to the core transcriptional regulatory circuitry of embryonic stem cells. Cell 134, 521–533 (2008).

40. Yi, F., Pereira, L. & Merrill, B. J. Tcf3 functions as a steady state limiter of transcriptional programs of mouse embryonic stem cell self renewal. 26, 1951–1960 (2008).

41. Liang, R. & Liu, Y. Tcf7l1 directly regulates cardiomyocyte differentiation in embryonic stem cells. Stem Cell Res. Ther. 9, 267 (2018).

42. Neagu, A. et al. In vitro capture and characterization of embryonic rosette-stage pluripotency between naive and primed states. Nat. Cell Biol. 22, 534–545 (2020).

43. Wu, C.-I. et al. Function of Wnt/β-catenin in counteracting Tcf3 repression through the Tcf3-β-catenin interaction. Development 139, 2118–2129 (2012).

44. Lackner, A. et al. Cooperative genetic networks drive embryonic stem cell transition from naïve to formative pluripotency. EMBO J. 9, 1–23 (2021).

45. Morgani, S., Nichols, J. & Hadjantonakis, A. The many faces of Pluripotency: in vitro adaptations of a continuum of in vivo states. BMC Dev. Biol. 17, (2017).

46. Plusa, B., Piliszek, A., Frankenberg, S., Artus, J. & Hadjantonakis, A. Distinct sequential cell behaviours direct primitive endoderm formation in the mouse blastocyst. Development 135, 3081–3091 (2008).

47. Morris, S. A. et al. Origin and formation of the first two distinct cell types of the inner cell mass in the mouse embryo. Proc. Natl. Acad. Sci. U. S. A. 107, 6364–6369 (2010).

48. Xenopoulos, P., Kang, M., Puliafito, A., DiTalia, S. & Hadjantonakis, A. K. Heterogeneities in Nanog expression drive stable commitment to pluripotency in the mouse blastocyst. Cell Rep. 10, 1508–1520 (2015).

49. de Jaime-Soguero, A., Abreu de Oliveira, W. A. & Lluis, F. The Pleiotropic Effects of the Canonical Wnt Pathway. Genes (Basel). 9, 93 (2018).

50. Mohammed, H. et al. Single-Cell Landscape of Transcriptional Heterogeneity and Cell Fate Decisions during Mouse Early Gastrulation. Cell Rep. 20, 1215–1228 (2017).

51. Nowotschin, S. et al. The emergent landscape of the mouse gut endoderm at single-cell resolution. Nature 569, 361–367 (2019).

52. Valenta, T., Hausmann, G. & Basler, K. The many faces and functions of b -catenin. EMBO J. 31, 2714–2736 (2012).

53. Tulac, S. et al. Identification, characterization, and regulation of the canonical Wnt signaling pathway in human endometrium. J. Clin. Endocrinol. Metab. 88, 3860–3866 (2003).

54. Mohamed, O. A. et al. Uterine Wnt/β-catenin signaling is required for implantation. Proc. Natl. Acad. Sci. U. S. A. 102, 8579–8584 (2005).

55. Hayashi, K., Ohta, H., Kurimoto, K., Aramaki, S. & Saitou, M. Reconstitution of the mouse germ cell specification pathway in culture by pluripotent stem cells. Cell 146, 519–532 (2011).

56. Franco, H. L. et al. WNT4 is a key regulator of normal postnatal uterine development and progesterone signaling during embryo implantation and decidualization in the mouse. FASEB J. 25, 1176–1187 (2011).

57. Wray, J., Kalkan, T., Gomez-lopez, S., Eckardt, D. & Cook, A. Inhibition of glycogen synthase kinase-3 alleviates Tcf3 repression of the pluripotency network and increases embryonic stem cell resistance to differentiation. Nat. Cell Biol. 13, 838–845 (2011).

58. ten Berge, D. et al. Embryonic stem cells require Wnt proteins to prevent differentiation to epiblast stem cells. Nat. Cell Biol. 13, 1070–1075 (2011).

59. Ryan, A. Q., Chan, C. J., Graner, F. & Hiiragi, T. Lumen Expansion Facilitates Epiblast-Primitive Endoderm Fate Specification during Mouse Blastocyst Formation. Dev. Cell 51, 684–697.e4 (2019).

60. Chatterjee, S. S. et al. Inhibition of β-catenin–TCF1 interaction delays differentiation of mouse embryonic stem cells. J. Cell Biol. 211, 39–51 (2015).

61. Pereira, L., Yi, F. & Merrill, B. J. Repression of Nanog Gene Transcription by Tcf3 Limits Embryonic Stem Cell Self-Renewal. Mol. Cell. Biol. 26, 7479–7491 (2006).

62. Lluis, F. et al. T-cell factor 3 (Tcf3) deletion increases somatic cell reprogramming by inducing epigenome modifications. Proc. Natl. Acad. Sci. U. S. A. 108, 11912–7 (2011).

63. Yi, F. et al. Opposing effects of Tcf3 and Tcf1 control Wnt stimulation of embryonic stem cell self-renewal. Nat. Cell Biol. 13, 762–770 (2011).

64. Merrill, B. J., Gat, U., DasGupta, R. & Fuchs, E. Tcf3 and Lef1 regulate lineage differentiation of multipotent stem cells in skin. Genes Dev. 15, 1688–1705 (2001).

65. Semrau, S. et al. Dynamics of lineage commitment revealed by single-cell transcriptomics of differentiating embryonic stem cells. Nat. Commun. 8, 1096 (2017).

66. Pruszak, J., Ludwig, W., Blak, A., Alavian, K. & Isacson, O. CD15, CD24, and CD29 define a surface biomarker code for neural lineage differentiation of stem cells. Stem Cells 27, 2928–2940 (2009).

67. Mitsui, K. et al. The Homeoprotein Nanog Is Required for Maintenance of Pluripotency in Mouse Epiblast and ES Cells. Cell 113, 631–642 (2003).

68. Ma, Z., Swigut, T., Valouev, A., Rada-Iglesias, A. & Wysocka, J. Sequence-specific regulator Prdm14 safeguards mouse ESCs from entering extraembryonic endoderm fates. 18, 120–127 (2011).

69. Nishiyama, A. et al. Uncovering Early Response of Gene Regulatory Networks in ESCs by Systematic Induction of Transcription Factors. Cell Stem Cell 5, 420–433 (2009).

70. Bencsik, R. et al. Improved transgene expression in doxycycline-inducible embryonic stem cells by repeated chemical selection or cell sorting. Stem Cell Res. 17, 228–234 (2016).

71. Brown, K. et al. A Comparative Analysis of Extra-Embryonic Endoderm Cell Lines. PLoS One 5, e12016 (2010).

72. Paca, A. et al. BMP signaling induces visceral endoderm differentiation of XEN cells and parietal endoderm. Dev. Biol. 361, 90–102 (2012).

73. Kunath, T. et al. Imprinted X-inactivation in extra-embryonic endoderm cell lines from mouse blastocysts. Development 132, 1649–61 (2005).

74. Artus, J. et al. BMP4 signaling directs primitive endoderm-derived XEN cells to an extraembryonic visceral endoderm identity. Dev. Biol. 361, 245–62 (2012).

75. Pfister, S., Steiner, K. A. & Tam, P. P. L. Gene expression pattern and progression of embryogenesis in the immediate post-implantation period of mouse development. Gene Expr. Patterns 7, 558–573 (2007).

76. Artus, J. et al. BMP4 signalling directs primitive endoderm-derived XEN cells to an extraembryonic visceral endoderm identity. Dev. Biol. 361, 245–262 (2013).

77. Frade, J. et al. Controlled ploidy reduction of pluripotent 4n cells generates 2n cells during mouse embryo development. Sci. Adv. 5, eaax4199 (2019).

78. Gerovska, D. & Araúzo-Bravo, M. J. Computational analysis of single-cell transcriptomics data elucidates the stabilization of Oct4 expression in the E3.25 mouse preimplantation embryo. Sci. Rep. 9, (2019).

79. Morgani, S. M. & Brickman, J. M. LIF supports primitive endoderm expansion during pre-implantation development. 3488–3499 (2015) doi:10.1242/dev.125021.

80. Kalkan, T. et al. Complementary Activity of ETV5, RBPJ, and TCF3 Drives Formative Transition from Naive Pluripotency. Cell Stem Cell 24, 785–801.e7 (2019).

81. Acampora, D., Di Giovannantonio, L. G. & Simeone, A. Otx2 is an intrinsic determinant of the embryonic stem cell state and is required for transition to a stable epiblast stem cell condition. Dev. Stem Cells 140, 43–55 (2013).

82. Zhang, J. et al. OTX2 restricts entry to the mouse germline. Nature 562, 595–599 (2018).

83. Yu, L. et al. Derivation of Intermediate Pluripotent Stem Cells Amenable to Primordial Germ Cell Specification. Stem Cell S1934-5909, 30541–5 (2021).

84. Han, H. et al. TRRUST v2: An expanded reference database of human and mouse transcriptional regulatory interactions. Nucleic Acids Res. 46, D380–D386 (2018).

85. Zhou, Y. et al. Metascape provides a biologist-oriented resource for the analysis of systems-level datasets. Nat. Commun. 10, 1523 (2019).

86. Cole, M. F., Johnstone, S. E., Newman, J. J., Kagey, M. H. & Young, R. A. Tcf3 is an integral component of the core regulatory circuitry of embryonic stem cells. Genes Dev. 746–755 (2008) doi:10.1101/gad.1642408.4.

87. Posfai, E. et al. Evaluating totipotency using criteria of increasing stringency. Nat. Cell Biol. 23, 49–60 (2021).

88. Schrode, N., Saiz, N., Di Talia, S. & Hadjantonakis, A. K. GATA6 levels modulate primitive endoderm cell fate choice and timing in the mouse blastocyst. Dev. Cell 29, 454–467 (2014).

89. Singh, A. M., Hamazaki, T., Hankowski, K. E. & Terada, N. A Heterogeneous Expression Pattern for Nanog in Embryonic Stem Cells. Stem Cells 25, 2534–2542 (2007).

90. Marks, H. et al. The Transcriptional and Epigenomic Foundations of Ground State Pluripotency. Cell 149, 590–604 (2012).

91. Kang, M., Garg, V. & Hadjantonakis, A. K. Lineage Establishment and Progression within the Inner Cell Mass of the Mouse Blastocyst Requires FGFR1 and FGFR2. Dev. Cell 41, 496–510.e5 (2017).

92. Kang, M., Piliszek, A., Artus, J. & Hadjantonakis, A. K. FGF4 is required for lineage restriction and salt-and-pepper distribution of primitive endoderm factors but not their initial expression in the mouse. Dev. 140, 267–279 (2013).

93. Bessonnard, S. et al. Gata6, Nanog and Erk signaling control cell fate in the inner cell mass through a tristable regulatory network. Development 141, 3637–3648 (2014).

94. Hamilton, W. B. & Brickman, J. M. Erk Signaling Suppresses Embryonic Stem Cell Self-Renewal to Specify Endoderm. Cell Rep. 9, 2056–2070 (2014).

95. Kemler, R. et al. Stabilization of β-catenin in the mouse zygote leads to premature epithelial-mesenchymal transition in the epiblast. Development 131, 5817–24 (2004).

96. Haegel, H. et al. Lack of beta-catenin affects mouse development at gastrulation. Development 121, 3529–37 (1995).

97. Chazaud, C. & Rossant, J. Disruption of early proximodistal patterning and AVE formation in Apc mutants. Development 133, 3379–3387 (2006).

98. Merrill, B. J. et al. Tcf3 : a transcriptional regulator of axis induction in the early embryo. (2004) doi:10.1242/dev.00935.

99. Harwood, B. N., Cross, S. K., Radford, E. E., Haac, B. E. & De Vries, W. N. Members of the WNT signaling pathways are widely expressed in mouse ovaries, oocytes, and cleavage stage embryos. Dev. Dyn. 237, 1099–1111 (2008).

100. Hayashi, K. et al. Wnt genes in the mouse uterus: Potential regulation of implantation. Biol. Reprod. 80, 989–1000 (2009).

101. Lyashenko, N. et al. Differential requirement for the dual functions of β-catenin in embryonic stem cell self-renewal and germ layer formation. Nat. Cell Biol. 13, 753–61 (2011).

102. Aulicino, F. et al. Canonical Wnt Pathway Controls mESC Self-Renewal Through Inhibition of Spontaneous Differentiation via β-Catenin/TCF/LEF Functions. Stem Cell Reports 15, 646–661 (2020).

103. Tam, W.-L. et al. T-Cell Factor 3 Regulates Embryonic Stem Cell Pluripotency and Self-Renewal by the Transcriptional Control of Multiple Lineage Pathways. Stem Cells 26, 2019–2031 (2008).

104. Morrison, G., Scognamiglio, R., Trumpp, A. & Smith, A. Convergence of cMyc and β-catenin on Tcf7l1 enables endoderm specification. EMBO J. 35, 356–368 (2016).

105. Mao, C. D. & Byers, S. W. Cell-context dependent TCF/LEF expression and function: Alternative tales of repression, de-repression and activation potentials. Crit. Rev. Eukaryot. Gene Expr. 21, 207–236 (2011).

106. Berasain, C. & Avila, M. A. Deciphering liver zonation: New insights into the β-catenin, Tcf4, and HNF4α triad. Hepatology 59, 2080–2082 (2014).

107. Moreira, S. et al. TCF7L1 and TCF7 differentially regulate specific mouse ES cell genes in response to GSK-3 inhibition. http://10.0.4.77/473801 (2018) doi:10.1101/473801.

108. Merrill, B. J. et al. Tcf3 : a transcriptional regulator of axis induction in the early embryo. Development 131, 263–274 (2004).

109. Liao, Y., Smyth, G. K. & Shi, W. FeatureCounts: An efficient general purpose program for assigning sequence reads to genomic features. Bioinformatics 30, 923–30 (2014).

110. Love, M. I., Huber, W. & Anders, S. Moderated estimation of fold change and dispersion for RNA-seq data with DESeq2. Genome Biol. 15, 550 (2014).

111. Reimand, J., Kull, M., Peterson, H., Hansen, J. & Vilo, J. g:Profiler—a web-based toolset for functional profiling of gene lists from large-scale experiments. Nucleic Acids Res. 35, W193–W200 (2007).

112. Robinson, J. T. et al. Integrative genomics viewer. Nat. Biotechnol. 29, 24–26 (2011).

113. Van De Sande, B. et al. A scalable SCENIC workflow for single-cell gene regulatory network analysis. Nat. Protoc. 15, 2247–2276 (2020).

114. Stuart, T. et al. Comprehensive Integration of Single-Cell Data. Cell 177, 1888–1902.e21 (2019).

115. Setty, M. et al. Characterization of cell fate probabilities in single-cell data with Palantir. Nat. Biotechnol. 37, 451–460 (2019).

116. De Leeneer, K. et al. Flexible, scalable, and efficient targeted resequencing on a benchtop sequencer for variant detection in clinical practice. Hum. Mutat. 36, 379–387 (2015).

117. Boel, A. et al. BATCH-GE: Batch analysis of Next-Generation Sequencing data for genome editing assessment. Sci. Rep. 6, 30330–30330 (2016).

